# SimpleMating: R-package for Prediction and Optimization of Breeding Crosses Using Genomic Selection

**DOI:** 10.1101/2024.05.24.595600

**Authors:** Marco Antônio Peixoto, Rodrigo Rampazo Amadeu, Leonardo Lopes Bhering, Luís Felipe V. Ferrão, Patrício R. Munoz, Márcio F. R. Resende

## Abstract

Selecting parents and crosses is a critical step for a successful breeding program. The ability to design crosses with high means that will maintain genetic variation in the population is the goal for long-term applications. Herein, we describe a new computational package for mate allocation in a breeding program. SimpleMating is a flexible and open-source R package originally designed to predict and optimize breeding crosses in crops with different reproductive systems and breeding designs. Divided into modules, SimpleMating first estimates the cross performance (criterion), such as mean parental average, cross total genetic value, and/or usefulness of a set of crosses. The second module implements an optimization algorithm to maximize a target criterion while minimizing next-generation inbreeding. The software is flexible, allowing users to specify the desired number of crosses, maximum/minimum number of crosses per parent, and the maximum value of the parent relationship for creating crosses. As an outcome, SimpleMating generates a mating plan from the target parental population using single or multi-trait criteria. As example, we implemented and tested SimpleMating in a simulated maize breeding program obtained through stochastic simulations. The crosses designed via SimpleMating showed large genetic mean over time (up to 22% more genetic gain than conventional genomic selection programs, with lower genetic diversity decrease over time), supporting the use of this tool, as well as the use of data-driven decisions in applied breeding programs.

## 1 INTRODUCTION

An essential step in animal and plant breeding programs is the selection of parents at the beginning of each crossing cycle. The crossing plan is achieved in plant breeding programs by forming combinations between superior parental lines, either within heterotic groups or at the base population. The strategy aims to ensure that the genetic mean of the next breeding cycle surpasses the one from the cycle where the parents are selected, resulting in genetic gains across generations. In this context, genomic selection has become a well-established selection tool in plant breeding programs, with examples increasing short-term genetic gain in various applications in different breeding programs (Fritsche-Neto et al., 2021; Wartha & Lorenz, 2021; Marinho et al., 2022). Genomic selection can increase the accuracy of the selection in the early stages, guaranteeing more certainty in truncation selection while enabling the rapid recycling of parents (Labroo & Rutkoski, 2022). However, to maintain higher gains over several selection cycles, the parental combinations must generate and retain reasonable levels of genetic variation (Wientjes et al., 2023). Unfortunately, genomic selection accelerates the loss of genetic diversity through breeding cycles compared with breeding strategies based on phenotypic truncation selection (Bančič et al., 2021; Sabadin et al., 2022). This loss is higher in traits with lower heritabilities. In this context, how could biometricians and breeders maintain performance and diversity in harmonic levels in the long term?

Over the last decades, several endeavors have been put together to increase genetic gain while maintaining or not depleting population genetic variation, particularly after the inclusion of genomic selection into breeding pipelines (Jannink, 2010; Daetwyler et al., 2015; Akdemir & Sánchez, 2016; Gorjanc & Hickey, 2018; Allier et al., 2019). The first attempt was reported by Meuwissen and Sonesson (1998), who suggested the concept of optimum contribution selection. The idea is to maximize the genetic value while limiting the rate of inbreeding (as a proxy for genetic variation). These estimates are used to estimate the optimal contributions of a set of parents to the next generation. The optimum contribution selection has served as a solid foundation for all subsequent contributions. For example, Jannink (2010) proposed to weigh allelic effects by their frequency and effect magnitude, aiming to preserve rare favorable alleles with significant effects on the genetic value. Another approach was the optimal haploid value, wherein individuals are selected by their predicted haploid value for the best doubled-haploids lines derived from the cross (Daetwyler et al., 2015). Using a similar motivation, other authors have focused on algorithms to optimize genetic gain while maintaining genetic diversity by setting a mating plan in breeding populations (Akdemir & Sánchez, 2016; Gorjanc & Hickey, 2018). In this latter case, crosses are optimized to maximize genetic mean while mate allocation is done to minimize inbreeding of mating plan (optimal cross selection).

Regardless of the methodology, selecting the best parents and combinations can be framed in the context of usefulness (Schnell & Utz, 1976). This concept combines genetic mean and genetic gain of a cross given a quantitative trait. The estimate of the genetic gain for a cross depends on the intensity of selection and the progeny variance. The estimation of progeny variance has been studied by several authors, who proposed different alternatives based on phenotypic distance, molecular markers distance in parental lines, and progeny simulation via genetic map information (Utz et al., 2001; Hung et al., 2012; Mohammadi et al., 2015). One approach that combines the use of molecular markers with population and parental linkage disequilibrium to estimate progeny variance has been shown to achieve superior results (Lehermeier et al., 2017b; a).

When selecting the superior parents, the breeding values of related individuals are generally ranked together, which can lead to more related individuals being mated at the beginning of each breeding cycle, increasing inbreeding (Falconer & Mackay, 1996). While the estimate of usefulness is conceptually an important approach to maximize genetic gain, breeders normally will also consider the expected level of inbreeding/coancestry from the resulting crosses before defining the mating plan. One approximation to achieve this goal is to combine the optimal cross-selection with a criterion to develop a set of breeding crosses. The optimization delivers a mating plan that maximizes a criterion (cross usefulness, mean parental average, or cross total genetic value) under constraints for expected relatedness, total number of crosses, and minimum/maximum number of crosses per parent. Long-term inbreeding rates are also reduced by keeping parental inbreeding/coancestry at low levels. Furthermore, avoiding mating relatives helps maintain short-term inbreeding rates at lower levels (Kinghorn, 2011). By controlling both short- and long-term inbreeding rates, we avoid depleting genetic variation and achieve higher long-term genetic gain.

In this study, we introduce an R package that can integrate cross-prediction and optimization using genomic data (Figure 1). The R package SimpleMating was designed with the following main objectives: (i) prediction of cross performance, such as mean parental average, cross total genetic value, and usefulness, and (ii) optimization of cross selection to increase a target criterion and restrict coancestry and/or the number of crosses per parent. The package can be used for models that assume additive or additive plus dominance effects. In addition, it does not require the DNA markers to be phased for usefulness prediction (even though it is accepted for specific scenarios). The package also allows cross-prediction using a single or several traits at a time. It is freely available on GitHub (https://github.com/Resende-Lab/SimpleMating). In the following, we describe the theory and technical implementation of the package, demonstrate how to use it, and evaluate its performance in a simulated maize breeding program. We end with a discussion on opportunities and future directions for the field.

**Figure 1.**
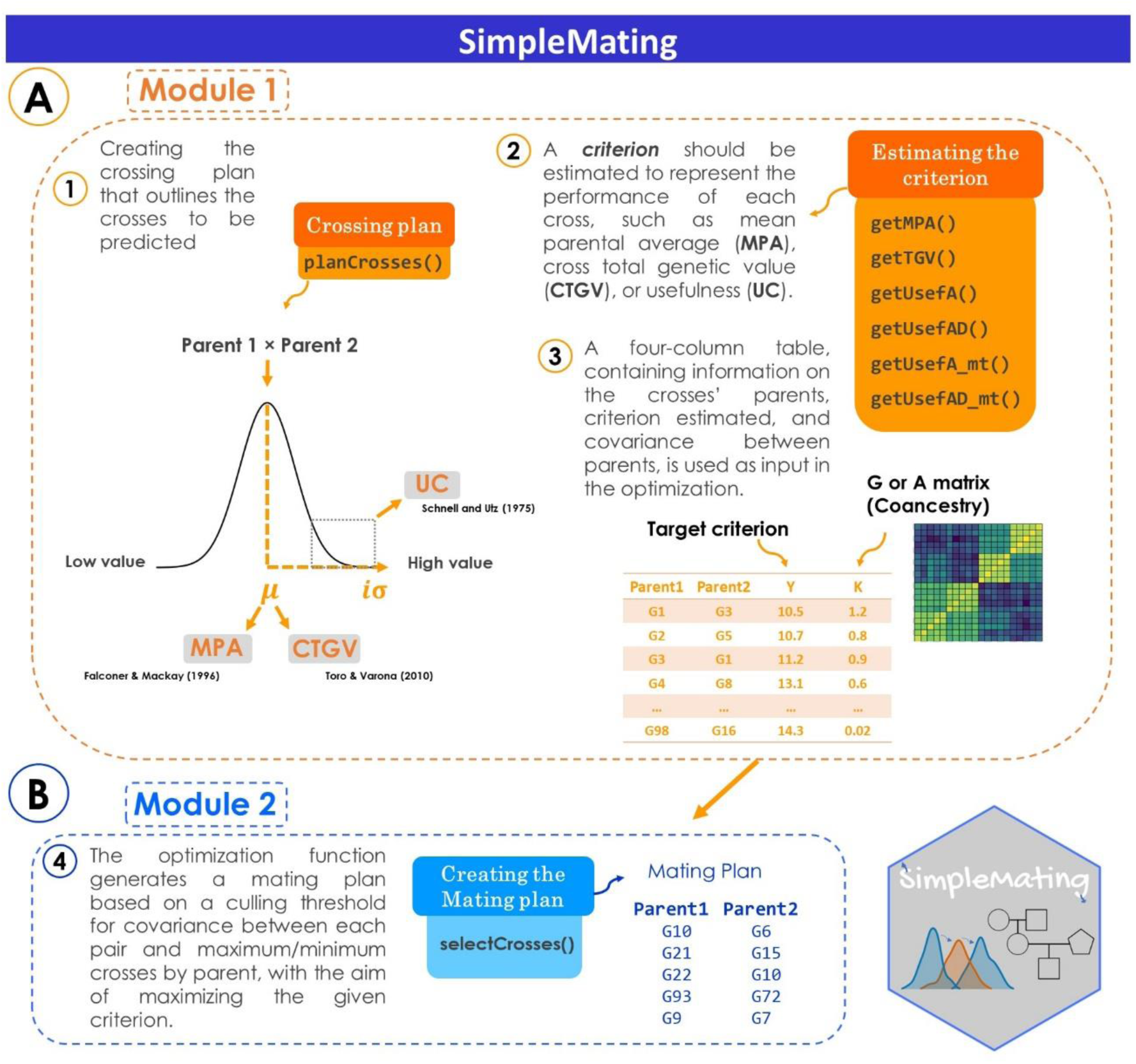
Workflow for the pipeline implemented in the SimpleMating package. Orange and blue colors represent the two main modules of the package: (A) estimation of cross-performance and (B) optimization. MPA: mean parental average, UC: usefulness, CTGV: cross-total genetic value. G: realized genomic relationship matrix. A: numerator relationship matrix.

## 2 USAGE AND APPLICATION

Here, the SimpleMating package functionality is described while connecting the functions with a real example. The user will find (i) the initial data input needed, (ii) a function to create the crossing plan, (iii) how to estimate the usefulness of a set of crosses, (iv) data input for the optimization, (v) the implementation of the optimization to retrieve a mating plan. Furthermore, the basic idea is to maximize a criterion with constraints for the rates of future inbreeding employing an optimization algorithm.

### 2.1 Data input

The initial step is data preparation. SimpleMating needs four inputs, but not all of them are required.

**1. Allele dosage matrix:** a matrix that contains the single nucleotide polymorphism (SNP) markers scores as allele dosage is required. It should include all potential candidates as parents. Here, the matrix is used to predict the contribution of each duo of parents to the future cross.

**2. Relationship matrix**: a matrix that could be a genomic-based relationship matrix (**G**), built up from an allele dosage matrix, or a pedigree-based relationship matrix (**A**) is required to connect the genotypes from the population. The package uses such information as a measure of coancestry to constrain inbreeding in the next generation, *i*.*e*., covariance from the G or A matrix between a pair of parents is twice the coancestry rate, which represents the inbreeding of the next generation for a set of crosses (Kinghorn, 2011).

**3. Genetic map/linkage disequilibrium**: a table with chromosome number, markers names, and position of the markers in the chromosomes (in centimorgan) (genetic map information). This is used to estimate the recombination matrix expected in the target population of parents. One alternative is to use the linkage disequilibrium between the markers from the population as a proxy to estimate the recombination among SNPs.

**4. Markers effects**: column vector (for one trait) or data frame (for more than one trait) containing the effect of each marker present in the allele dosage matrix. Both additive and dominance marker effects should be supplied in different arguments for a trait with additive and dominance control. For several traits, the user may supply marker effects for all traits from the same set of SNPs.

### 2.2 Installation and loading

After organizing the input files required, the next step is to install the package SimpleMating and load it. The user may install it directly from GitHub using the ‘devtools’ package as follows:

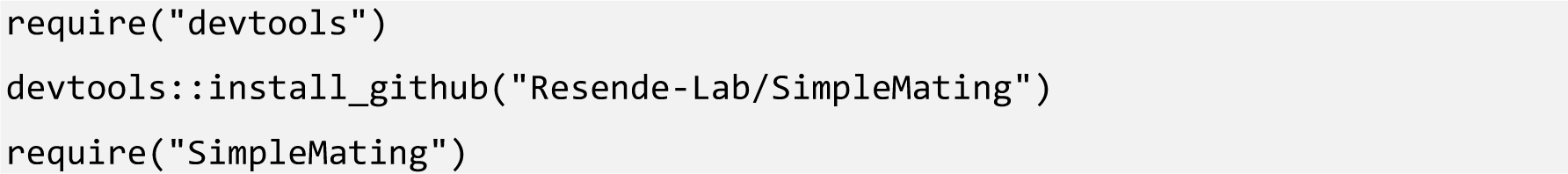

### 2.3 Planning crosses

The first module starts with planning the crosses. The possible crosses are created using the list of possible parents. The function `planCross()` can be used to generate a data frame with all crosses to be optimized. The user can also come up with its own crossing plan without using this function. In this function, the argument *TargetPop* represents the names of potential parents to be mated. The argument *MateDesign* represents the type of crosses that should be generated. The options are: ‘*full*’: indicates that all parents should be crossed with all parents, considering reciprocal crosses. ‘*full_p*’: indicates that all parents should be crossed with all parents, considering self of parents, and reciprocal crosses. ‘*half’*: indicates that neither reciprocal nor parents are included in the crossing plan. ‘*half_p*’: indicates that all parents could be crossed with all parents, considering self of parents but not reciprocal crosses.

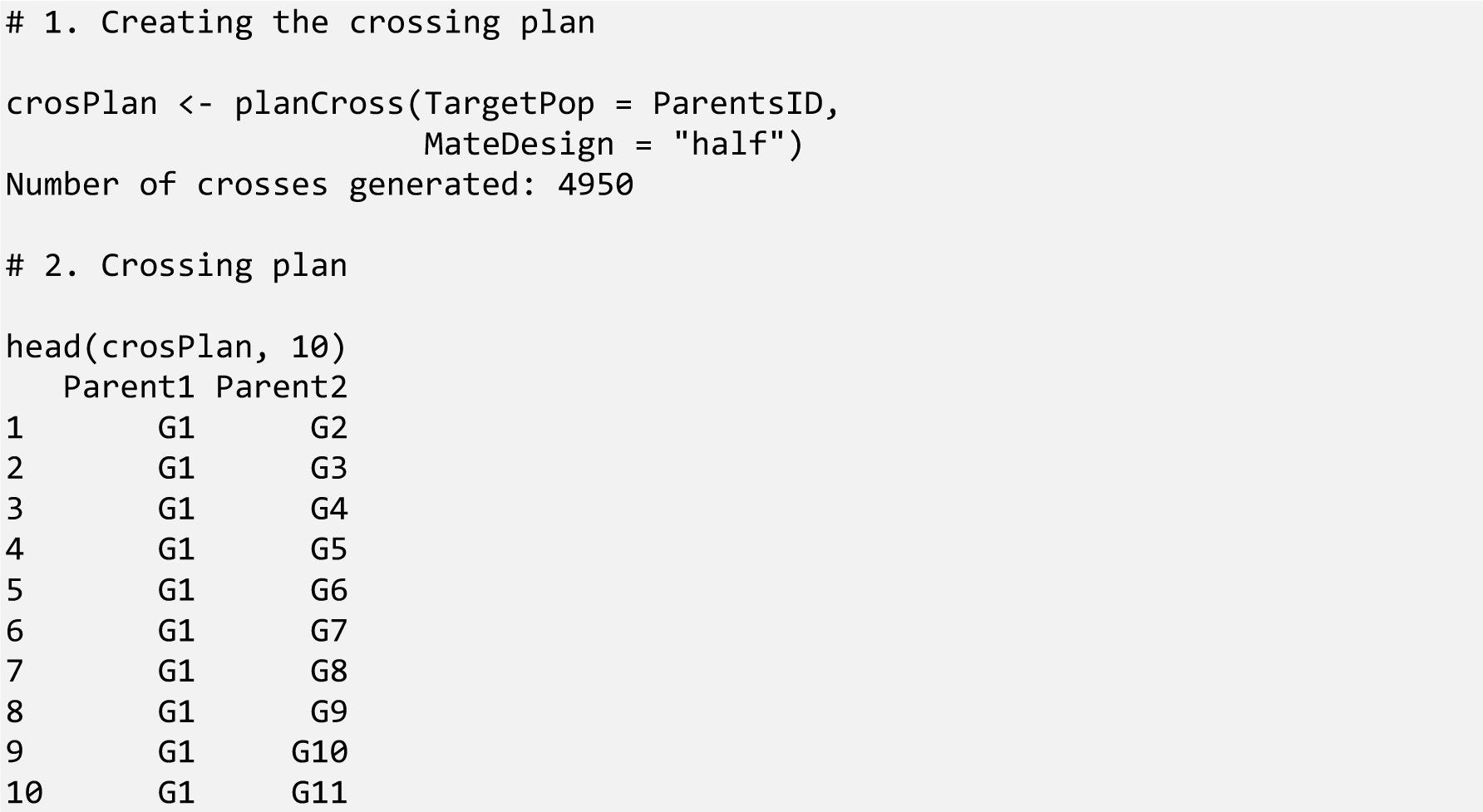

In addition to the usage described above, other useful arguments in this function are *Indiv2keep* and *TargetPop2*. The argument *Indiv2keep* asks for a list of all individuals kept after a thin-out process and can be used with the `relateThinning()` function (see section 2.6 for more information). When argument *TargetPop2* is used, the individuals from population one (*TargetPop*) are crossed solely with the individuals from population two (*TargetPop2*). It resembles the North Carolina II mating design (Comstock and Robinson, 1952). This provides the possibility of generating crosses between two populations, which might be beneficial in different cases such as hybrid make-ups, sex and/or logistic limitations, etc.

### 2.4 Cross-criterion estimation

After simply generating the potential crosses for the parental population (or across populations), the next step is to predict a criterion to be maximized based on the parents’ performance. SimpleMating gives the possibility to predict the mean parental average, cross total genetic value, or the estimation of usefulness, depending on the focus and input data. In the first case, assuming that only the BLUP of the parents is known (together with a pedigree-based relationship matrix for optimization – Module 2), the mean parental average (*μ_ebv_*) can be estimated for each cross in the crossing plan by using the function `getMPA()`, giving as output a data frame ready to be used as input in the optimization. Multiple trait scenarios can be used by setting weights for each trait.

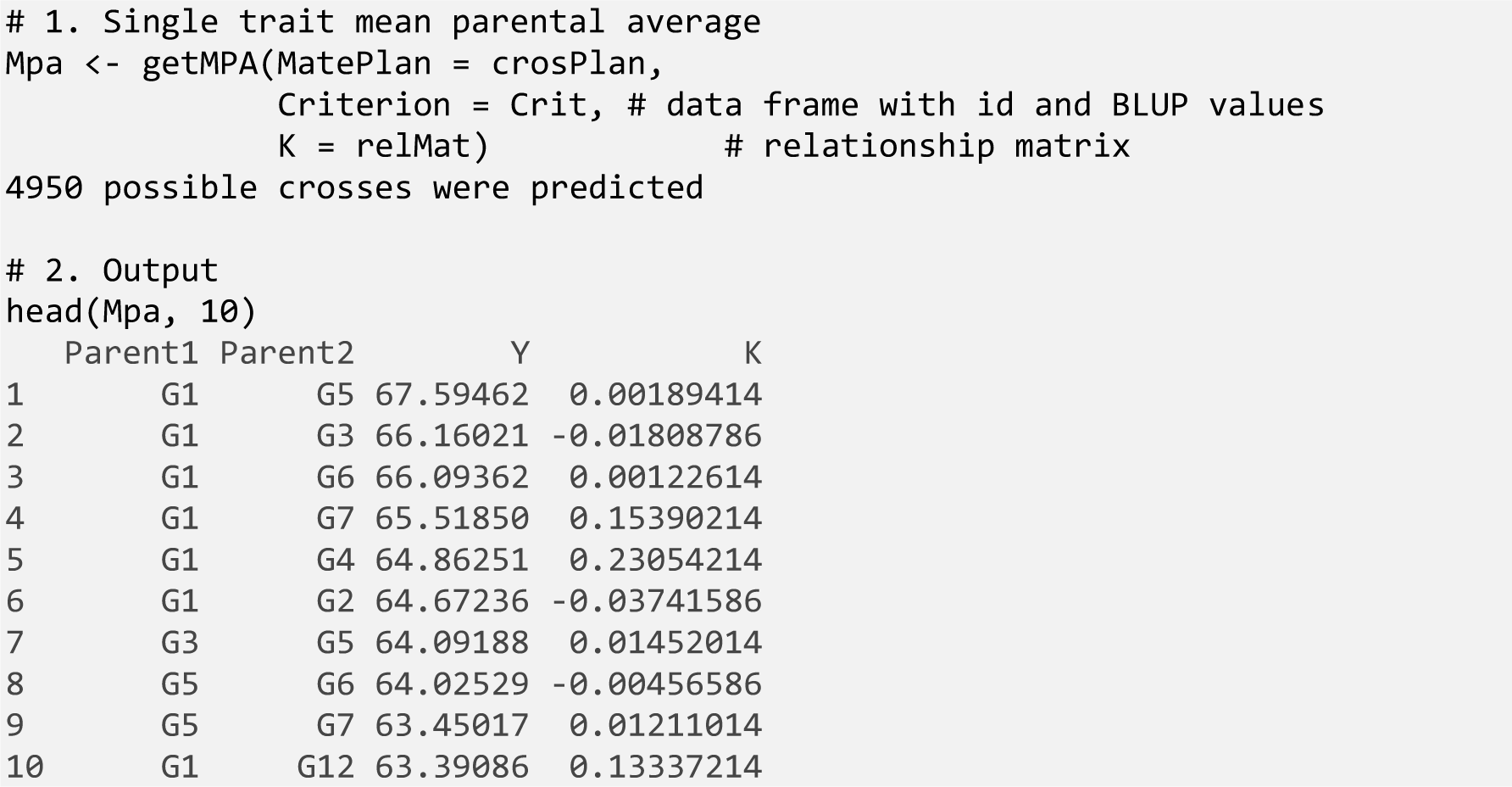

The second possibility is to use the function `getTGV()` to estimate the cross-total genetic value (for traits controlled by additive plus dominance effects) (Falconer & Mackay, 1996). The cross-total genetic value has been shown to improve genetic gain in the long run, especially for clonally propagated species (Werner et al., 2023). In this way, the average heterosis from a biparental cross (captured by dominance effects) in the F1 is capitalized in the parental selection. The following equations are used for the estimation (Falconer & Mackay, 1996; Toro & Varona, 2010):

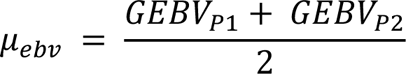

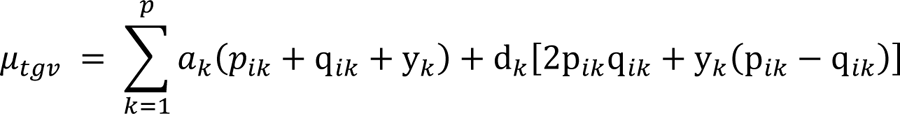

where *GEBV* represents the genomic *ebv* for each parent (being another criterion that defines a parent performance also accepted, such as BLUPs), *a* and *d* represent the additive and dominance effects of the markers, respectively, *p* and *q* represent the allele frequency for a marker, and *y* represents the difference in allele frequency between populations (parents) involved in a cross. Multiple trait scenarios can be used by setting weights for each trait.

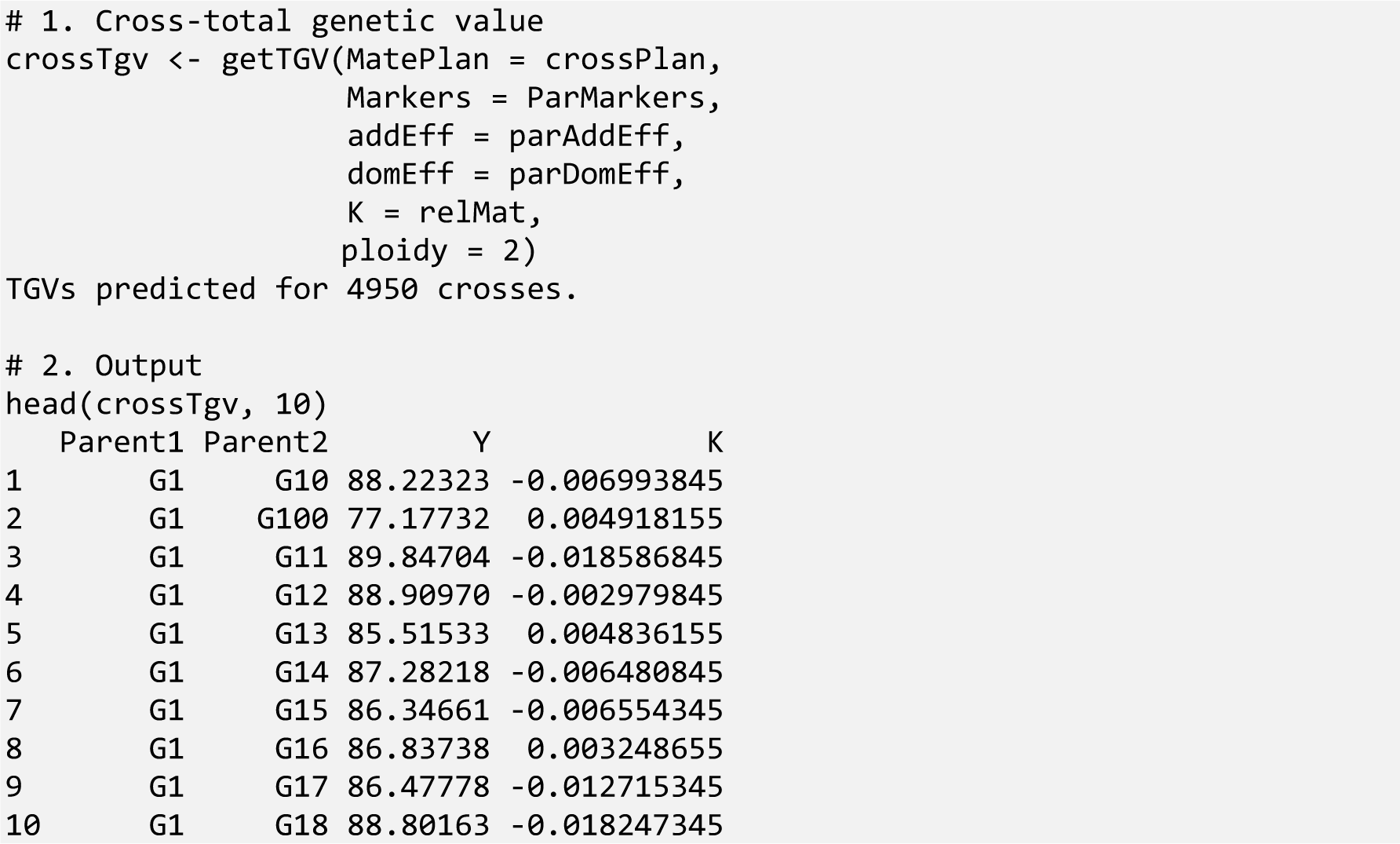

The last possibility in the first module is the prediction of the usefulness of the target traits/traits. Four functions can be used to select potential crosses based on their usefulness: one trait with additive control (*getUsefA*), one trait with additive and dominance control (*getUsefAD*), more than one trait with additive control (*getUsefA_mt*), and more than one trait with additive and dominance control (*getUsefAD_mt*). The usefulness criterion (UC) combines the concepts of genetic mean (*μ*) and genetic gain (Δ*G*) and seeks to maximize the performance of the next generation. Its implementation has proven to return higher gains than the selection of combinations based on only truncation selection (Lehermeier et al., 2017b; Allier et al., 2019). The following equation describes UC:

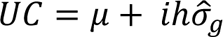

where μ is the mean of a target cross (genetic mean), *i* represents the intensity of selection, *h* is the squared root of the heritability (assumed to be 1), and 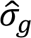 represents the progeny genetic standard deviation within a family for a target cross (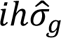 term stands for genetic gain - Δ*G*). In the abovementioned equation, selection intensity (*i*) represents the mean of deviations from the population mean. For the functions considering only additive effects, usefulness is estimated considering the mean parental average (*μ_ebv_*) as the mean term using the additive markers effects information and the dosage matrix. In contrast, for those considering additive and dominance effects, the mean estimated is the cross total genetic value (*μ_tgv_*). With this, we ensure that the usefulness captured dominance effects in the mean and the variance for usefulness. A thorough description of the calculation and implementation of the formulas for the usefulness estimates is given in Supplemental Material S1.

The cross-variance estimation is based on recombination frequencies of the parental population and the linkage disequilibrium covariance between loci of the parents for both additive and dominance variance (Bonk et al., 2016; Lehermeier et al., 2017b). In this approach, for traits with only additive gene actions, the functions accept doubled haploids (DH) and recombinant inbred lines (RIL) as populations from where the parents can be derived (single and multi-trait cases). As only homozygous individuals are expected in the population (either recessive or dominant homozygous), there is no need for phased haplotypes in this situation. However, for cross-targeting traits controlled by additive and dominance gene actions (single and multi-trait cases), the cross-variance estimation includes the need to know the phase of the markers from the set of parents. This information is useful for estimating linkage disequilibrium covariance between loci of a target cross (Bonk et al., 2016; Lehermeier et al., 2017b; Wolfe et al., 2021). As an alternative, we implemented in SimpleMating an equation to estimate linkage disequilibrium covariance between loci based on non-phased haplotypes (detailed in Supplemental Material S1). Further details and examples for all functions are presented in the package’ vignette.

Here, we described the implementation of one trait controlled by only additive effects (using *getUsefA*). The inputs are the allele dosage matrix (*Markers*), the additive effects for each marker (*addEff*), the relationship matrix (*K*, only used for setting up the output for the optimization), the genetic map for all markers (*Map.In*), the proportion of individuals to be selected (*propSel*), and crossing plan before estimated (*MatePlan*).

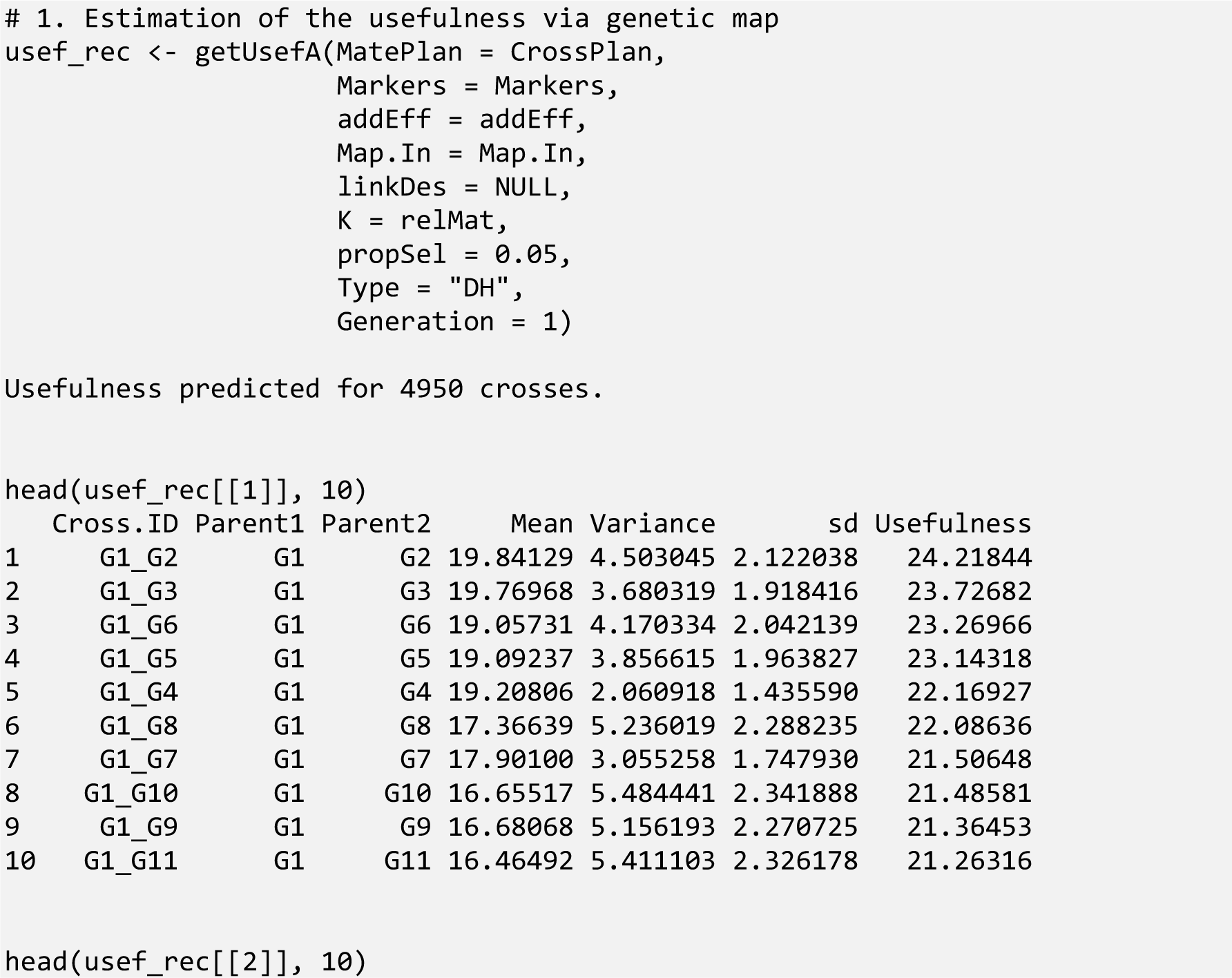
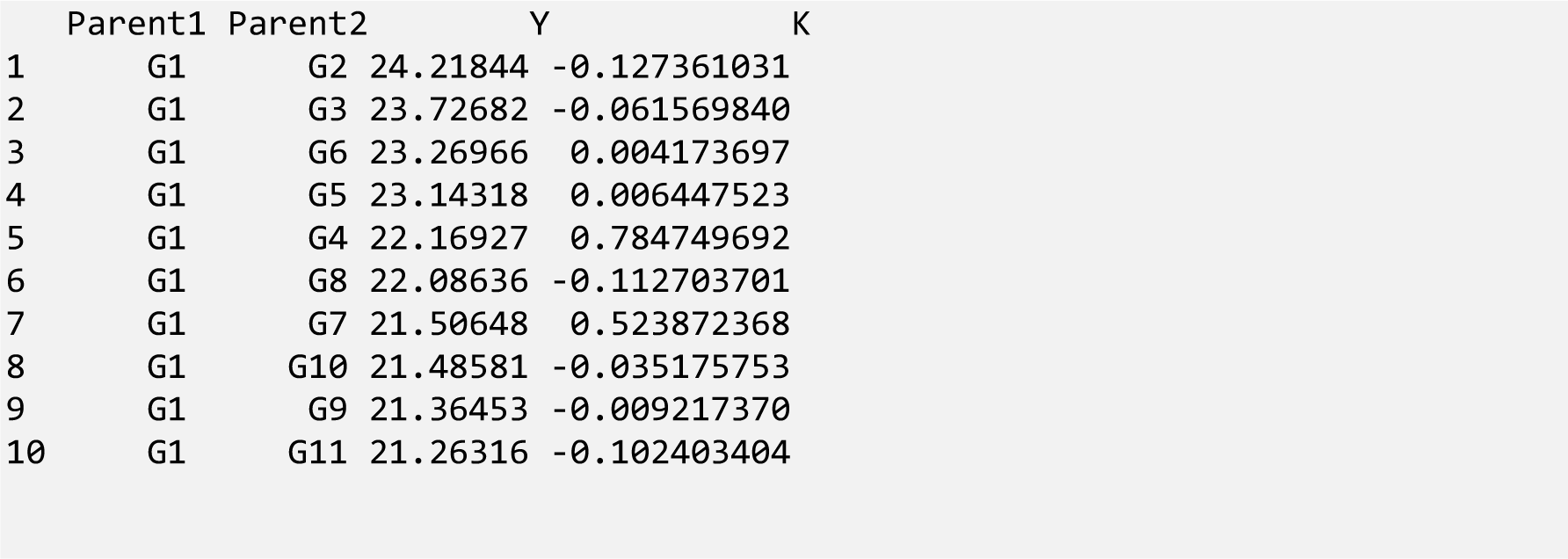

One alternative to the use of the genetic map (sometimes not simple to estimate for the target population) is to estimate the linkage disequilibrium (LD) matrix among markers and use it in the argument *linkDes*. The matrix will be used to construct a block diagonal matrix (number of blocks equals the number of chromosomes) to capture the recombination pattern in the marker data. It is worth noting that 1-LD is a proxy for recombination for a given base pair. In this case, the user should add the information of chromosome and marker name (chromosome + marker Id) for speeding up the process instead of the genetic map in the argument *Map.In*.

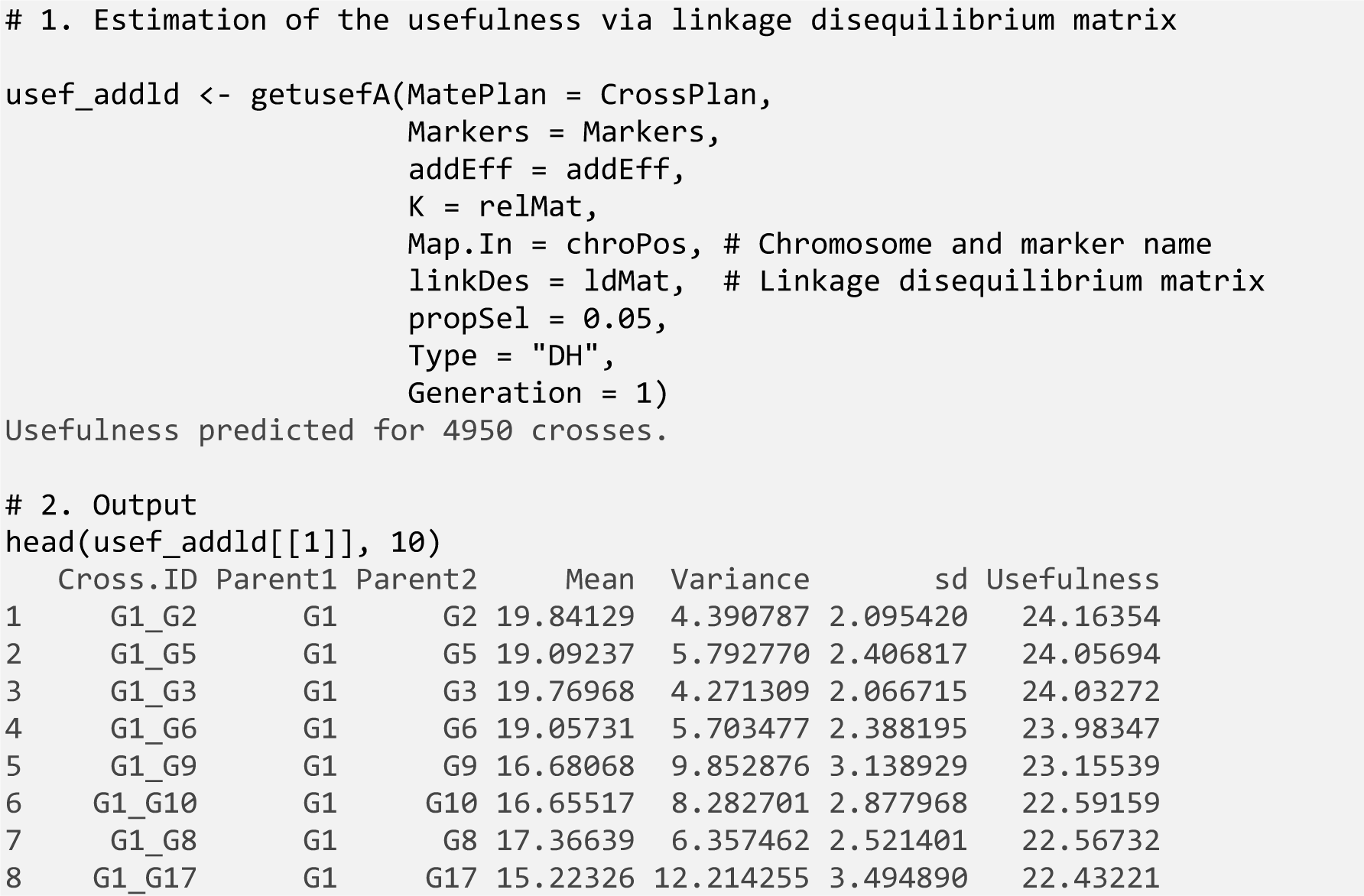
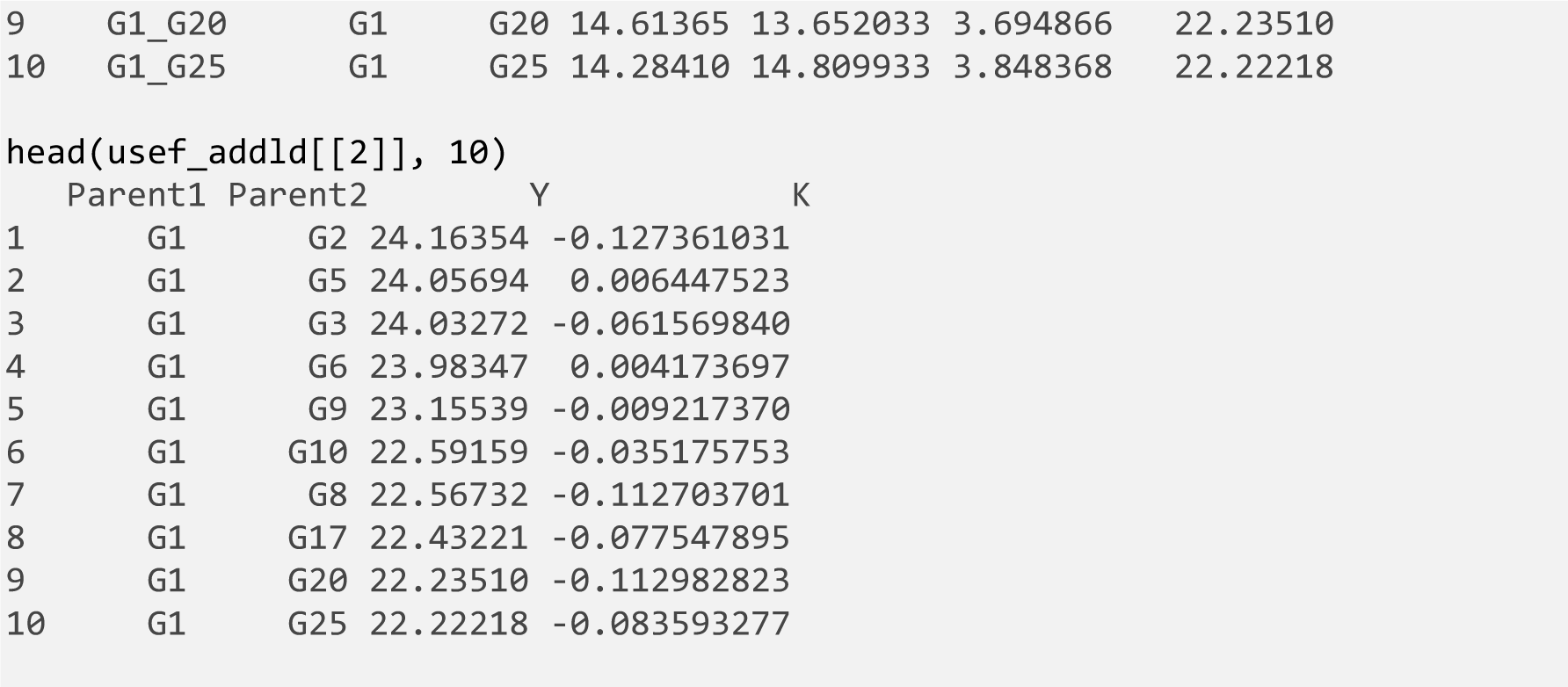

The result from the mean parental average (`getMPA()`) and cross total genetic value (`getTGV()`) functions is a table ready to be the optimization function input. The result is a two-item list object for the usefulness of predictions’ functions. The first position of the list contains cross id (Cross.ID), parents (Parent1 and Parent2), progeny mean (Mean), progeny variance (Variance), standard deviation (sd, progeny variance’ squared root), and usefulness for the trait (Usefulness) and in the second the input for the optimization function.

When we ranked the crosses in the example above (1. Estimation of the usefulness via genetic map) by the crosses’ mean and by the crosses’ usefulness, the cross pairs G1 × G7 and G1 × G8 switched rank positions. This has an implication when all crosses are ranked together and shows that the chosen ones may be different using the different criteria for selection (Figure 2), even though the cross usefulness and cross mean have a similar trend (Supplemental Material S2- Figure 2). This may cause an impact in the long run (more about this in the example below). In addition, the use of the linkage disequilibrium information for the recombination matrix estimation instead of the genetic map seems to have a similar pattern, varying in the magnitude of the variance (Supplemental Material S2 - Figure 2).

**Figure 2.**
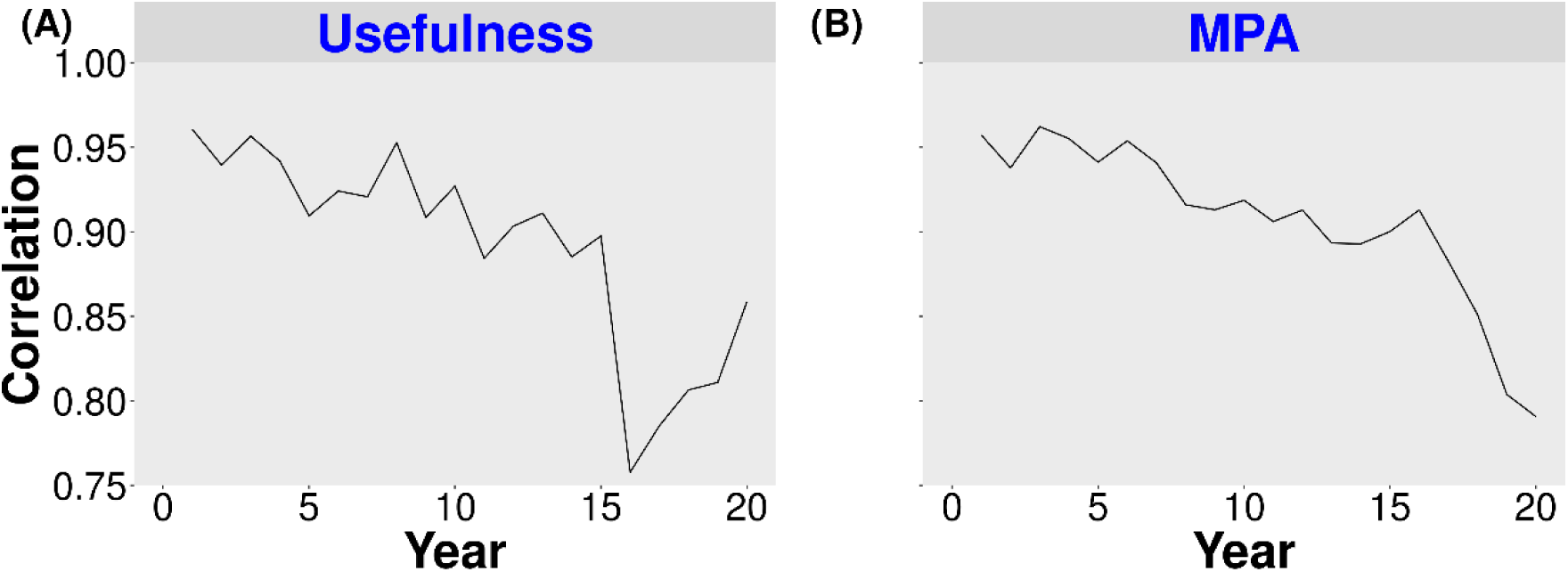
Correlation between usefulness and mean parental average (MPA) over 20 years of simulation. Selection of crosses at the beginning of each breeding cycle was made based on usefulness criterion (A) and mean parental average (B).

### 2.5 Running the optimization

After building the input file, the user may implement the optimization via the **`selectCrosses`**() function. The optimization algorithm from this function works with a four-column data frame as input in the argument *data*, as given: Parent1 and Parent2 columns represent the parents of the target cross, Y represents the criterion to be maximized, and K is a column with the covariance values. Column Y, representing the *criterion,* can contain the estimated mean parental average, cross-total genetic value, or cross-usefulness. In most cases, we expect the user to use the direct output of the functions described (topic 2.4) in the optimization.

Moreover, the function requires the total number of crosses that should be in the mating plan (*n.cross*), maximum (*max.cross*) and minimum contribution (*min.cross*) of each parent, maximum number of crosses from the input data that should be analyzed (*i*.*e*., rows in the dataset presented in *data* argument) to search (*max.cross.to.search*), and the culling pairwise (*culling.pairwise.k*) based on the covariance among individuals from the relationship matrix, where the restriction by relatedness is set. This argument is used to trim every cross with a covariance value higher than this value. All of the remaining crosses will be ranked based on the criterion (Y column), and the top ones will be selected based on *n.cross* input. After, from those selected crosses, a restriction is made by the number of crosses each parent is part of (*max.cross* and *min.cross* arguments). After, the process is repeated until the mating plan contains a set of crosses that follows all arguments in the function.

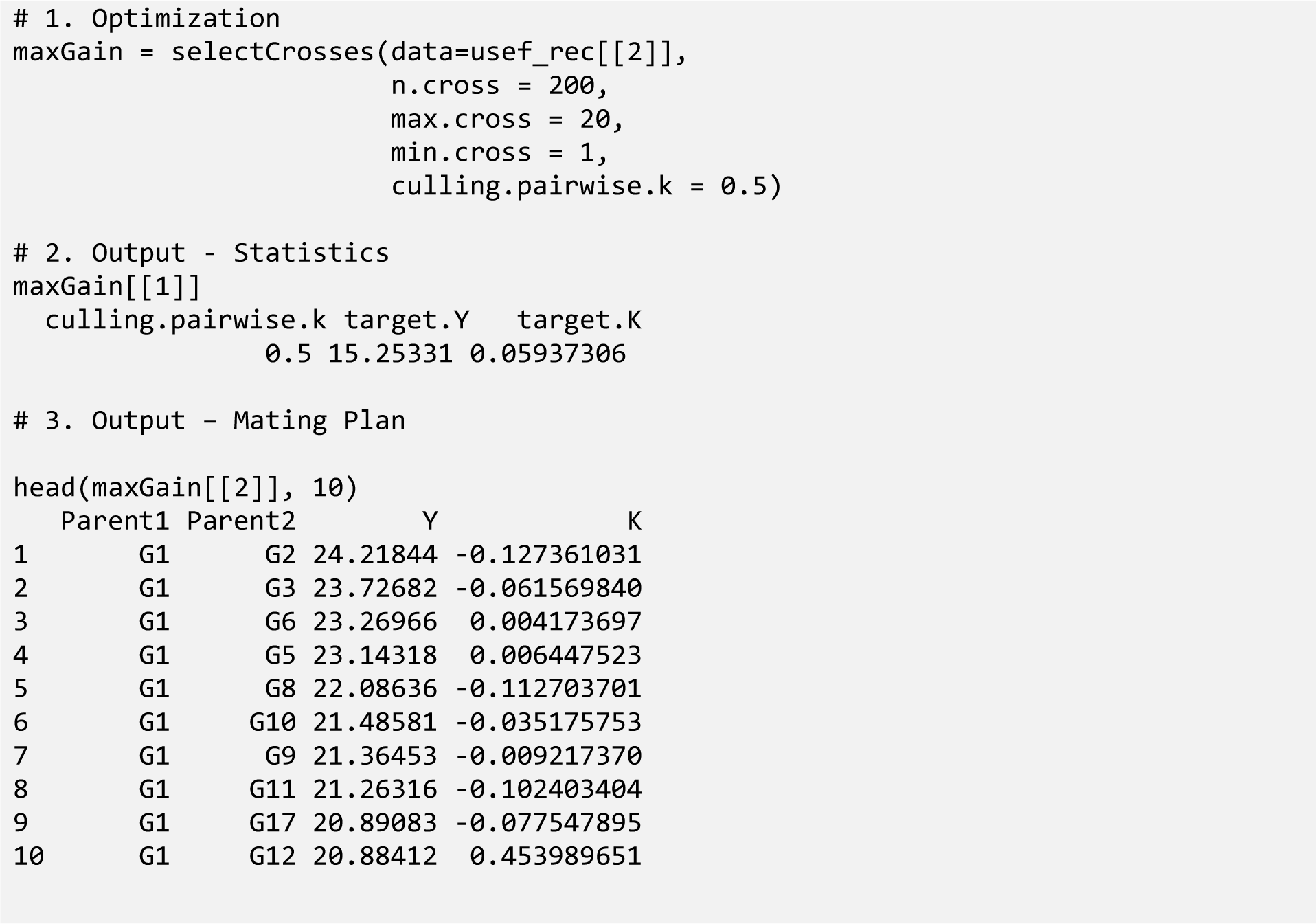

After the optimization, the output is a three-fold object, in which the user can assess (i) the optimization parameters targeted by the optimization (*culling.pairwise.k*, *target.Y*, and *target.k*), (ii) the mating plan accordingly, (iii) a graph showing the selected crosses in respect with the total crosses comparing the criterion used (Y) and the relatedness set via *K* argument (Figure 3). The mating plant is built based on the parameters set in the optimization. For instance, changes in the number of crosses that each parent can be part of and the culling for covariance restriction have an impact on the mating plan (see Figure 3), and the user can change them for targeting the breeding program’s specificities. A set of detailed examples with large datasets for different traits is presented in the vignette and is available at https://github.com/Resende-Lab/SimpleMating.

**Figure 3.**
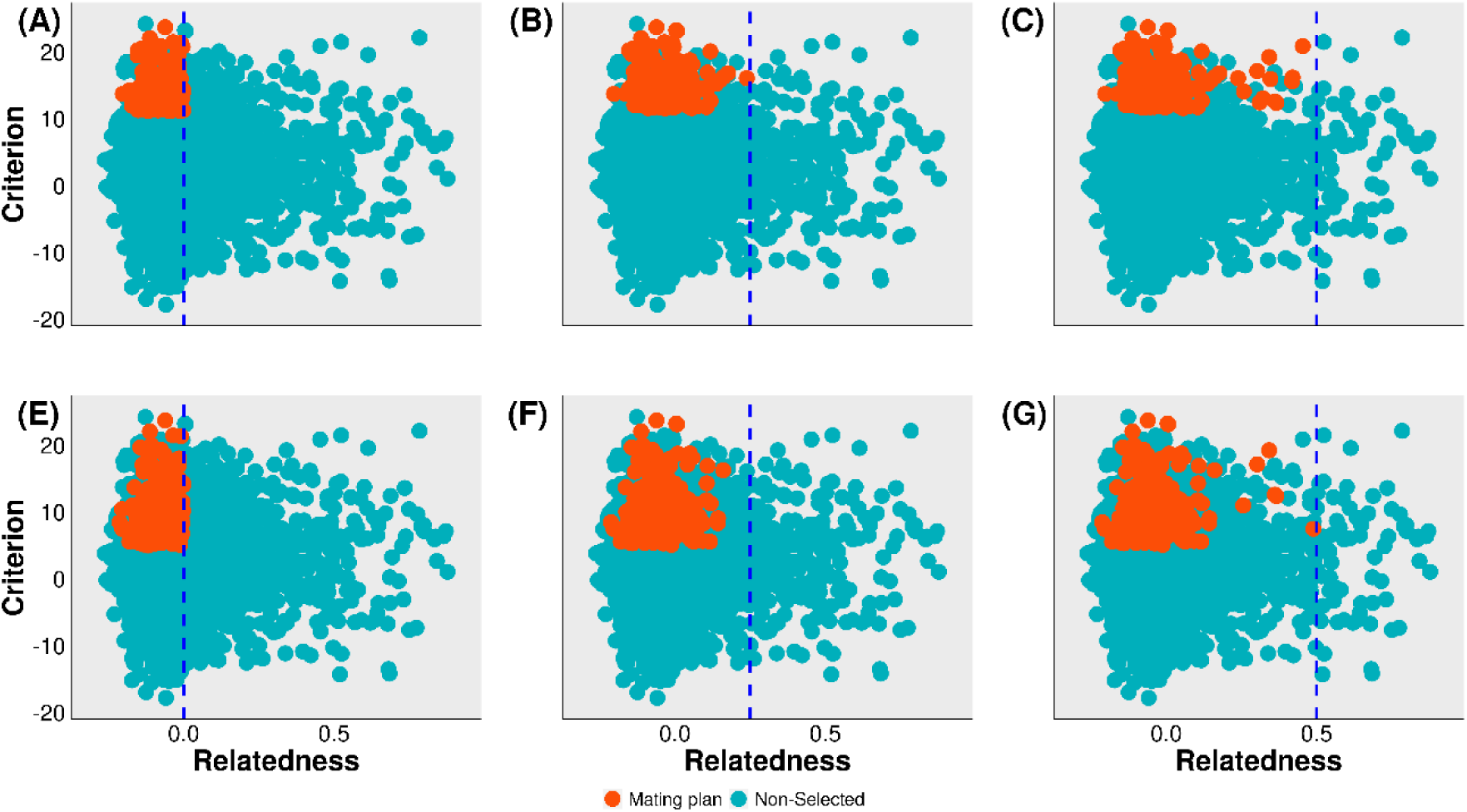
Mating plan (red dots) as an outcome from the selectCrosses() function varying the *culling.pairwise.k* (dotted blue line) and *max.cross* arguments. The first argument was set to 0 (Covariance 0, A and E), 0.25 (Covariance 0.25, B and F), and 0.5 (Covariance 0.5, C and G). The total number of crosses that each parent can be part of (*max.cross*) was set to 20 (A, B, and C) and 5 (E, F, and G) and the *min.cross* argument was set to 1 in all of them. Only the crosses’ pair with a value lower than the covariance set via *culling.pairwise.k* are considered candidates for crosses.

### 2.7. Speeding up the process

The estimation of the cross criterion is part of the first module of the package. In the case of usefulness computation, the variance estimation could be time-consuming, especially when the number of markers and/or number of crosses to estimate is high. SimpleMating works internally in a by-chromosome approach, and only segregating markers between pairs of parents is used to estimate the cross variance (for usefulness). These approaches are used to speed up the process. We reported some benchmarks to use SimpleMating in a couple of scenarios (different number of crosses and number of markers), reporting the total time combining usefulness estimation and optimization (Figure 4). It is worth noting that the time may vary from population to population once the number of segregating markers may differ between pair of parents.

**Figure 4.**
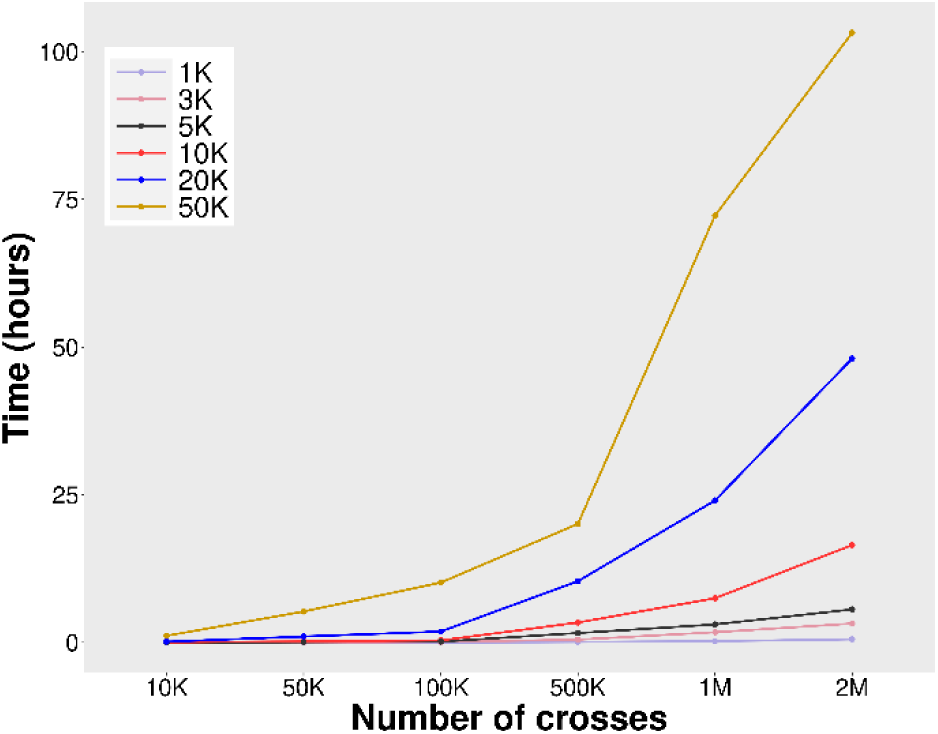
Time to predict usefulness (module 1) and to optimize (module 2) for a set of parents using SimpleMating. The time for the optimization (module 2) was small/disregarded. Each line represents one set of markers varying from 1 thousand to 50 thousand markers. The number of crosses varied from 10 thousand to 2 million crosses predicted. A set of 100 crosses were generated in the mating plan. For all runs, HiperGator (3.0 GHz Cores) was used, with 6 CPUs and 10GB of memory.

For a large number of individuals and crosses, the time for getting a mating plan can be significant (more than 100 hours for 50 thousand SNPs and 2 million crosses predicted). To decrease the time to get the mating plan, individuals can be trimmed beforehand. For such, the function `relateThinning()` can be employed to thin out some individuals considered potential parents, even though it remains an optional step. By reducing the number of selected candidates through the removal of closely related individuals (*i*.*e*., those with high covariance values in the given relationship matrix), the time required for predicting usefulness and cross-optimization decreases. As a result, the efficiency of the process is enhanced. A detailed explanation of the function is provided in Supplemental Material S2 – Figure A and an example is given in the main vignette.

This function implements an algorithm to select the top individuals based on a relatedness threshold defined by the user. For instance, if *max*.*per*.*cluster* = 2 and *threshold* = 0.5, the algorithm will identify all groups of individuals that shared more than 0.5 in the relationship matrix and select two individuals from the group with the highest criterion to be used as potential parents. In addition to these arguments, the function requires the relationship matrix (*K* argument) between the parents (*n* × *n*, being *n* the number of candidates to parents). Furthermore, the *criterion* argument should include information about the individual’s performance for the target trait (*n* × 1). Here, the user can set the estimated breeding value of the individual, the total genetic value, or the phenotypic values of the candidates. Then, as a result, a list of individuals to keep is generated and can be used directly in the argument *Indiv2keep* in the function that creates the crossing plan (`planCross()`).

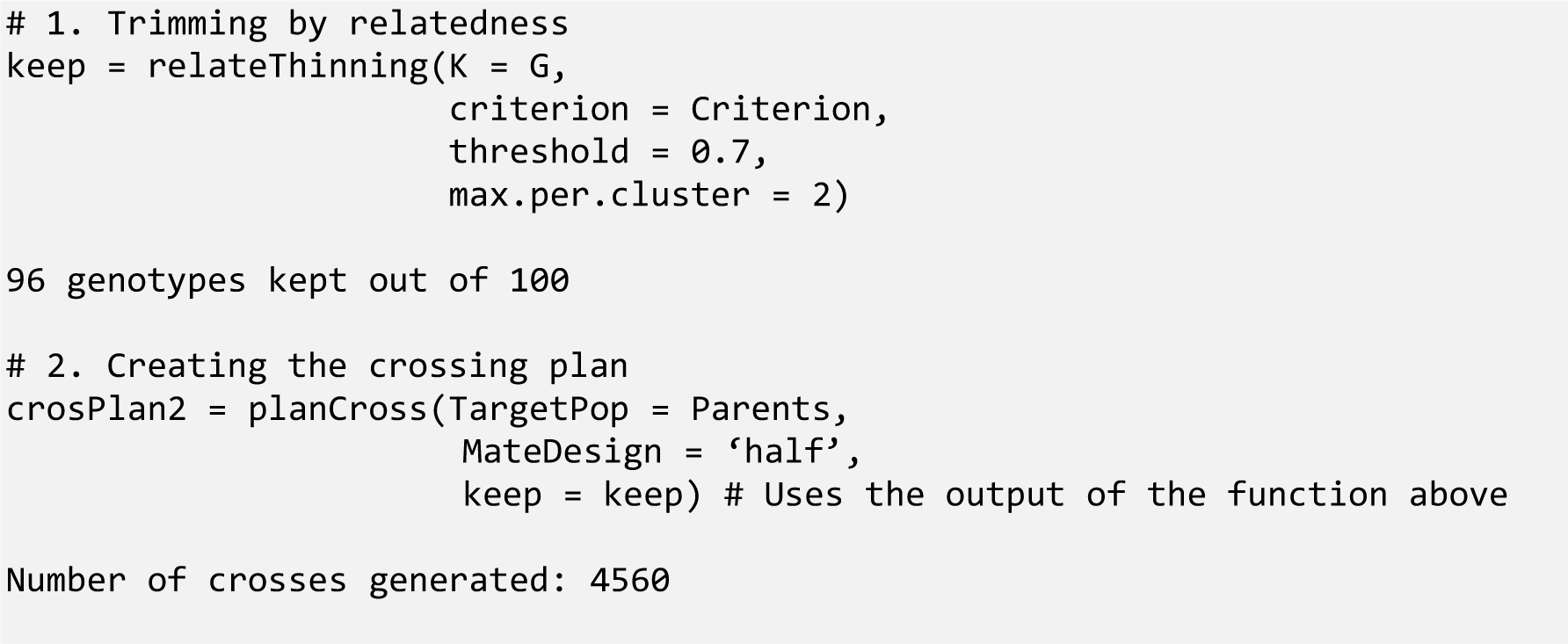

In this example, with a thinning of four individuals, we kept 96 individuals out of 100. In such case, we will predict usefulness or mean parental average/total genetic value for 4560 crosses instead of 4950 crosses (the number of crosses resulted from matching 100 individuals using *MateDesign = ‘half’*).

## 3 DEMONSTRATION

A demonstration of SimpleMating is presented through a set of stochastic simulations. A reciprocal recurrent selection maize breeding program was simulated (described by Powell et al., 2020). The program simulated the use of doubled haploid technology to generate homogenous lines from F_1_ populations (Figure 5). All tested scenarios were generated over a 20-year breeding program. A 15-year burn-in phase was implemented so each scenario would start at the same point. A burn-in phase represents the past years of the breeding program, while the 20 years represent the upcoming years, where the strategies will be evaluated and compared. Each scenario was replicated 20 times.

**Figure 5.**
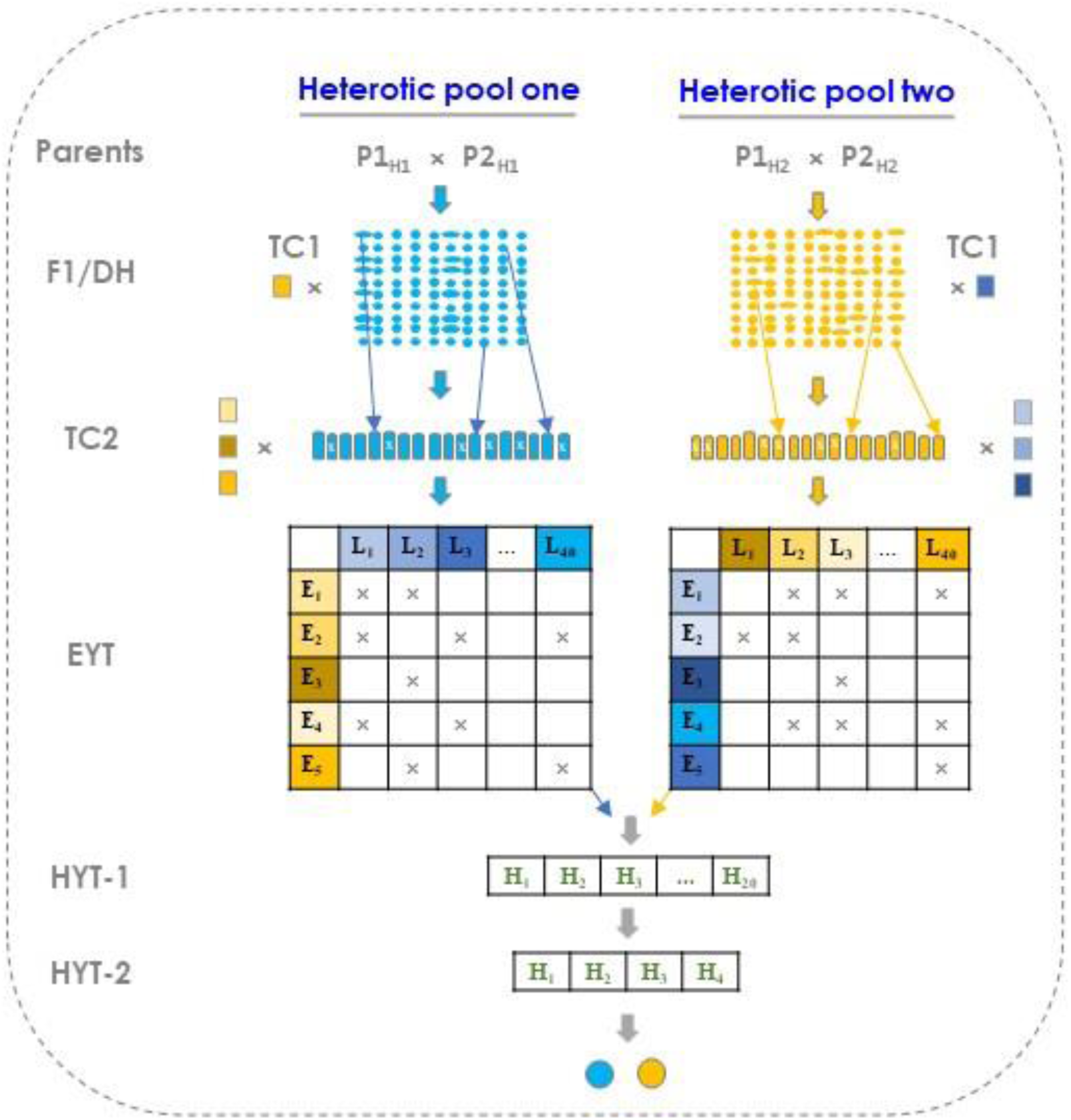
Schematical representation of a simulated maize breeding program. Two heterotic pools were simulated. In the first stage, doubled haploids were generated from F_1_ lines. Two sets of test crosses were implemented (TC1 and TC2). The hybrids were created by crossing lines from one heterotic group (colors orange and blue) with elite lines that came from the other heterotic group. In the end, the selection was made based on the hybrid performance in three different years, and the two best hybrids from each heterotic group were released as hybrid varieties. HYT: hybrid yield trial (one and two) YET: elite yield trial.

### 3.1 Base genome and trait simulations

The first step was the implementation of a base genome containing 10 chromosome pairs via the Markovian Coalescent Simulator (Chen et al., 2009), available in AlphaSimR (Gaynor et al., 2020; R Development Core Team, 2022). It used the standard ‘*Maize’,* which brings all genetic/genomic characteristics of the crop into the simulation (Hickey et al., 2014; Powell et al., 2020 for more details). A total of 300 quantitative trait nucleotides were distributed evenly per pair of chromosomes (3000 total). A split of the base genome into two different core genomes was simulated 100 generations ago. That split mimics the two heterotic groups structured in maize (*i*.*e*., flint and dent groups), which are used as two different groups for a reciprocal recurrent selection program. The biological model for this set of simulations was an additive, dominance, and genotype-by-environment interaction model. For the implementation of genomic selection model in the breeding program, 1000 single nucleotide polymorphisms (SNPs) were randomly allocated across each chromosome pair, representing a total of 10000 SNPs for each genotype.

One trait was simulated with a genetic mean of 70, a genetic variance of 20, a genotype-by-environment of 40, and a residual variance of 270. The dominance effect was calculated according to the dominance degree of the trait, with a mean of 0.92 and a variance of 0.3. These values assumed for the mean and variance of the dominance degree were chosen to follow the historical levels that roughly represent the heterosis in field corn for yield trait (Troyer & Wellin, 2009; Powell et al., 2020).

### 3.2 Scenarios

Five scenarios were simulated for comparison. Before it started, the burn-in period (to mimic the development of maize breeding in the past years) was set using the reciprocal recurrent selection as the breeding strategy. This strategy starts by creating a new round of 40 F_1_s (40 crosses randomly assigned) from 40 elite parents in the program, followed by successive cycles of reciprocal recurrent selection, for each heterotic pool. All selections in the burn-in phase were based on observed phenotypic values, and 15 years of cycles were performed. After the burn-in phase, the five different scenarios were run.

The first one (*PS* scenario) used the same pipeline as the burn-in period, *i*.*e*., parents came from a truncation phenotypic selection followed by 40 random crosses into the reciprocal recurrent selection program. The second scenario was the implementation of genomic selection in the pipeline, where the parents were 40 individuals from DH stage were selected by their estimated breeding value performance at the begging of each cycle, whereas the mating cross was randomly allocated (*GS* scenario) for each heterotic pool.

The third scenario was outlined by the usefulness criterion estimation for each possible combination (crosses) out of the DH stage that contains 1000 individuals. The target usefulness was the criterion maximized, whereas the number of crosses per parent and covariance values was restricted to create a mating plan with 40 crosses per cycle via the SimpleMating package (hereafter called *OCS* scenario). The maximum/minimum number of crosses of each parent was set to 3 and 1, respectively. Also, the *culling.pairwise.k* was to be set to 0. A genetic map was used to estimate usefulness for each cross combination coming from the 1000 individuals, for each heterotic pool.

The next scenario, same implementation was used than *OCS* scenario, but as target criterion we used the mean parental average (*MeanPAve* scenario) for each possible cross combination, estimated by SimpleMating. Restrictions for the number of crosses of each parent and inbreeding rates were implemented. For this case, the maximum/minimum number of crosses of each parent was set to 3 and 1, respectively. Also, the *culling.pairwise.k* parameter was set to 0.5.

In the last scenario, was implemented using the same pipeline as the *PS* scenario, but the crosses were assigned via SimpleMating. For this scenario (so-called *OCSPed*), a pedigree-based relationship matrix (A matrix) was used to generate estimated breeding values for all individuals from the DH stage. The estimated breeding value of each individual was used to estimate the mean parental average using SimpleMating for each potential cross. This information was used as input in the optimization function. The maximum/minimum number of crosses of each parent was set to 3 and 1, respectively, while a *culling.pairwise.k* was set up to 1.75.

In the scenarios *GS*, *MeanPAve* and *OCS*, marker effects were estimated by fitting a random regression best linear unbiased prediction (RRBLUP) model (Meuwissen et al. 2001) using the RRBLUP function implemented in AlphaSimR. The genomic model used in all scenarios was calibrated in a slide-window approach, were the three recently years (starting at year 13 of burn-in years) were used to calibrate the model with the phenotypes and genotypes coming from the general combining ability of testcrosses one and two. One genomic model for each heterotic pool was generated.

In the *OCSPed*, the estimated breeding values were generated using BLUPF90 program (Misztal et al., 2018), by running a pedigree-based model (Pocrnic et al., 2023). The pedigree information was tracked and used for the genomic model since the first year of the burn-in period. The numerator relationship matrix (A matrix) was calculated using all information on pedigree available for individuals from F_1_ population and all doubled haploids.

The performance of each scenario was compared by measuring the parents’ parameters. The genetic gain was an average of the genetic values, that were scaled to have mean zero at year 15 (last year of burn-in period). We measured the genetic diversity with genetic standard deviation and genic standard deviation (Gorjanc et al., 2018). The genetic standard deviation was estimated as the standard deviation of the standardized genetic values. The genic standard deviation was estimated as the square root of the variance of genetic values assuming no linkage between the causal loci, directly obtained from AlphaSimR parental’s population (Gorjanc et al., 2018; Lara et al., 2022; Pocrnic et al., 2023). All scripts for the implementation of the simulated scenarios and to plot and analyze the results can be found at: https://github.com/Resende-Lab/MaizeOCS-SimpleMating.

## 4. RESULTS

Here, the use and applicability of SimpleMating were compared within a maize reciprocal recurrent selection program utilizing 20 years of simulation with 20 simulation replicates. In each cycle, 40 parents were selected and randomly mated in the benchmark scenarios (*PS* and *GS*), while in the other scenarios, SimpleMating assigned the crosses (*MeanPAve*, *OCSped,* and *OCS*). The results are presented in Figure 6.

**Figure 6.**
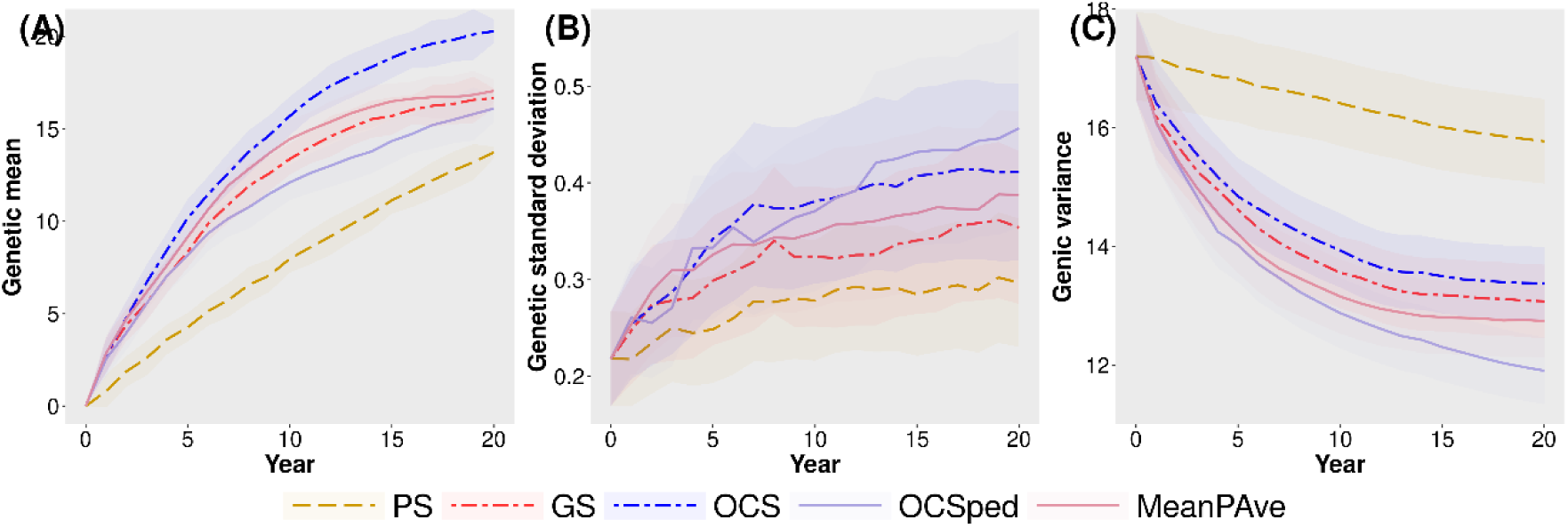
Genetic mean (A), genetic standard deviation (B), and genic variance (C) over time for five scenarios of a simulated maize breeding program. The scenarios are plotted as the average of the genetic value of the parents at each cycle, with 20 replicates. The shading around the line represents the standard error of the mean. *PS*: phenotypic selection scenario with random crosses assigned. *GS*: genomic selection with random crosses assigned; *OCSPed*: phenotypic selection and optimization of crosses using SimpleMating on mean parental average as criterion together with inbreeding restriction based on the A matrix from the pedigree. *OCS* genomic selection and optimal cross selection using SimpleMating package on usefulness as the target criterion together with covariance restriction. *MeanPAve*: genomic selection and optimization of crosses on mean parental average as the criterion through SimpleMating together with inbreeding restriction.

The population mean performance over a 20-year simulation period showed that the *PS* scenario resulted in a lower long-term genetic gain compared to other scenarios (*GS*: 21.43%, *MeanPAve*: 24.24%, *OCS*: 47.80%, and *OCSPed*: 17.31% of genetic gain higher than PS scenario). Among the scenarios that used genomic selection, *MeanPAve* slightly outperformed *GS* in the long-term result (2.31% higher), even though in the first few years of simulation the *MeanPAve* scenario showed a large gain compared with *GS* scenario. The *OCS* scenario was the one with higher performance (21.71% higher than GS at year 20). In addition, optimal cross selection used in scenarios *OCS* and *OCSped* conserved the most genetic diversity (genetic standard deviation), while the drop in genic variance was smaller compared with *GS* scenario for *OCS* scenario.

These results suggest that SimpleMating leverages the genetic gain in all scenarios where it was used, but the use of cross-usefulness is also important to keep long-term gain high. Further research should target other scenarios such as trait heritabilities, the influence of QTL number controlling the traits, marker panel size, and a larger number of traits.

## 5. DISCUSSION

The SimpleMating package is a computational and open-source tool designed for optimizing cross allocation, with the central goal of a target criterion and constrains inbreeding rates simultaneously. Written in the R language, which is widely popular in the fields of biology, statistics, and genetic data analysis, SimpleMating emerges as a valuable tool to make breeding decisions more data-oriented and maximize gains in the long-term. As R packages have been considered in the routine data analyses of quantitative genetics, SimpleMating was originally designed to easily connect with this framework in a way that regular outputs from any quantitative genetic analyses could be used directly in the software for cross allocation.

SimpleMating can be utilized in a wide range of breeding programs and various situations. For instance, it is particularly valuable for breeding programs that focus on multiple traits simultaneously, which is expected to be the most important. SimpleMating enables the estimation of the usefulness criterion for a group of traits, facilitating optimization based on the represented usefulness index. Furthermore, the ability to estimate and incorporate non-additive effects, such as dominance effects, can leverage the mating allocation for several crops where non-additive effects are important (Toro & Varona, 2010; Varona et al., 2018; Werner et al., 2023).

For breeding programs that do not employ genomic selection in their routine, crosses can be optimized based on pedigree records. To this end, the use of BLUP (via mean parental average) as the sole criterion for optimization and the A matrix for inbreeding rate constraints is highly encouraged. Our results demonstrated that utilizing robust pedigree information to guide parental crosses leads to significantly higher genetic gain compared to the *PS* scenario. Such a scenario can be particularly beneficial for small breeding programs, as it generates valuable information for data-driven decisions.

The assumptions made for the prediction of cross variance for the algorithms here implemented were for diploid-like species. However, for polyploid crops, such as potatoes, tomatoes, sugar cane, and sugar beet, among others, the use of SimpleMating is also advisable. Even though the prediction of cross variance and, consequently, the usefulness is not implemented, the use of the cross total genetic value together with inbreeding constraint is possible and is recommended. Werner et al. (2023) demonstrated the advantages of cross-total genetic value, and a similar pattern has been shown for polyploidy crops via simulation (results not shown). Conducting more analysis encompassing this example exceeds this study’s scope. Nevertheless, it is crucial to underscore the significance of such an implication.

## 6. CONCLUSION

Here, the SimpleMating package is described as a method for predicting crosses and optimizing genetic mean and inbreeding in breeding programs. The program enables both animal and plant breeding programs to be more optimal and facilitates new research opportunities.

## ACKNOWLEDGMENTS

This work was supported by USDA-NIFA 2018-51181-28419, USDA-NIFA 2019-05410, and USDA-NIFA 2022-51181-38333 to M.F.R.R.). We also thank the Brazilian government for the financial support from the National Council for Scientific and Technological Development (CNPq) and the Coordination for the Improvement of Higher Education Personnel (CAPES) through the CAPES-PrInt scholarship. This study was financed in part by the CAPES - Finance Code 001.

## SUPPLEMENTAL MATERIAL

**Supplemental Material S1:** Demonstration of genotypic variance-covariance of DH/RILs lines and the inclusion of non-additive effects for outbreed species derived from progenies of a biparental cross.

**Supplemental Material S2:** Figures describing functions and outputs.

## Abbreviations

BLUP: best linear unbiased prediction
GS: genomic selection
LD: linkage disequilibrium
MeanPAve: mean parental average
SNP: single-nucleotide polymorphism
OCS: optimal cross selection
PS: phenotypic selection.

## REFERENCES

Akdemir, D., & Sánchez, J.I. (2016). Efficient breeding by genomic mating. Frontiers in Genetics, 7, 1–12. 10.3389/fgene.2016.00210

Allier, A., Moreau, L., Charcosset, A., Teyssèdre, S., & Lehermeier, C. (2019). Usefulness criterion and post-selection parental contributions in multi-parental crosses: Application to polygenic trait introgression. G3: Genes, Genomes, Genetics, 9, 1469–1479. 10.1534/g3.119.400129

Bančič, J., Werner, C.R., Gaynor, R.C., Gorjanc, G., Odeny, D.A., Ojulong, H.F., Dawson, I.K., Hoad, S.P., & Hickey, J.M. (2021). Modeling Illustrates That Genomic Selection Provides New Opportunities for Intercrop Breeding. Frontiers in Plant Science, 12. 10.3389/fpls.2021.605172

Bonk, S., Reichelt, M., Teuscher, F., Segelke, D., & Reinsch, N. (2016). Mendelian sampling covariability of marker effects and genetic values. Genetics Selection Evolution, 48, 1–11. 10.1186/s12711-016-0214-0

Chen, G.K., Marjoram, P., & Wall, J.D. (2009). Fast and flexible simulation of DNA sequence data. Genome Research, 19, 136–142. 10.1101/gr.083634.108

Daetwyler, H.D., Hayden, M.J., Spangenberg, G.C., & Hayes, B.J. (2015). Selection on Optimal Haploid Value Increases Genetic Gain and Preserves More Genetic Diversity. Genetics, 200, 1341–1348. 10.1534/genetics.115.178038

Falconer, D.S., & Mackay, T.F.C. (1996). Introduction to quantitative genetics. Harlow, Essex, UK: Longmans Green, 3, 280

Fritsche-Neto, R., Galli, G., Borges, K.L.R., Costa-Neto, G., Alves, F.C., Sabadin, F., Lyra, D.H., Morais, P.P.P., Andrade, L.R.B., Granato, I., & Crossa, J. (2021). Optimizing Genomic-Enabled Prediction in Small-Scale Maize Hybrid Breeding Programs: A Roadmap Review. Frontiers in Plant Science, 12, 1–16. 10.3389/fpls.2021.658267

Gaynor, R.C., Gorjanc, G., & Hickey, J. (2020). AlphaSimR: An R-package for Breeding Program Simulations. G3 Genes|Genomes|Genetics, 1–21. 10.1101/2020.08.10.245167

Gorjanc, G., Gaynor, R.C., & Hickey, J.M. (2018). Optimal cross selection for long-term genetic gain in two-part programs with rapid recurrent genomic selection. Theoretical and Applied Genetics, 131, 1953–1966. 10.1007/s00122-018-3125-3

Gorjanc, G., & Hickey, J.M. (2018). AlphaMate : a program for optimizing selection, maintenance of diversity and mate allocation in breeding programs. Bioinformatics, 34, 3408–3411. 10.1093/bioinformatics/bty375

Hickey, J.M., Dreisigacker, S., Crossa, J., Hearne, S., Babu, R., Prasanna, B.M., Grondona, M., Zambelli, A., Windhausen, V.S., Mathews, K., & Gorjanc, G. (2014). Evaluation of genomic selection training population designs and genotyping strategies in plant breeding programs using simulation. Crop Science, 54, 1476–1488. 10.2135/cropsci2013.03.0195

Hung, H.Y., Browne, C., Guill, K., Coles, N., Eller, M., Garcia, A., Lepak, N., Melia-Hancock, S., Oropeza-Rosas, M., & Salvo, S. (2012). The relationship between parental genetic or phenotypic divergence and progeny variation in the maize nested association mapping population. Heredity, 108, 490–499

Jannink, J. (2010). Dynamics of long-term genomic selection. *Genetics*, Selection and Evolution, 42, 1–11

Kinghorn, B.P. (2011). An algorithm for efficient constrained mate selection. Genetics Selection Evolution, 43, 1–9

Labroo, M.R., & Rutkoski, J.E. (2022). New cycle, same old mistakesOverlapping vs discrete generations in long-term recurrent selection. BMC Genomics, 22, 1:15. 10.1101/2021.10.12.464059

Lara, L.A.C., Pocrnic, I., Oliveira, T.P., Gaynor, R.C., & Gorjanc, G. (2022). Temporal and genomic analysis of additive genetic variance in breeding programmes. Heredity, 128, 21–32. 10.1038/s41437-021-00485-y

Lehermeier, C., de los Campos, G., Wimmer, V., & Schon, C.-C. (2017)(a). Genomic variance estimates: With or without disequilibrium covariances?. Journal of Animal Breeding and Genetics, 134, 232–241. 10.1111/jbg.12268

Lehermeier, C., Teyssèdre, S., & Schön, C.C. (2017)(b). Genetic gain increases by applying the usefulness criterion with improved variance prediction in selection of crosses. Genetics, 207, 1651–1661. 10.1534/genetics.117.300403

Marinho, C.D., Coelho, I.F., Peixoto, M.A., Carvalho Júnior, G.A., & Resende Jr, M.F.R. (2022). Genomic selection as a tool for maize cultivars development. Revista Brasileira de Milho e Sorgo, 21

Meuwissen, T.H.E., & Sonesson, A.K. (1998). Maximizing the response of selection with a predefined rate of inbreeding: overlapping generations. Journal of Animal Science, 76, 2575–2583

Misztal, I., Lourenco, D., Aguilar, I., Legarra, A., & Vitezica, Z. (2018). Manual for BLUPF90 family of programs

Mohammadi, M., Tiede, T., & Smith, K.P. (2015). PopVar: A Genome-Wide Procedure for Predicting Genetic Variance and Correlated Response in Biparental Breeding Populations. Crop Science, 55, 2068–2077. 10.2135/cropsci2015.01.0030

Pocrnic, I., Obšteter, J., Gaynor, R.C., Wolc, A., & Gorjanc, G. (2023). Assessment of long-term trends in genetic mean and variance after the introduction of genomic selection in layers: a simulation study. Frontiers in Genetics, 14, 1–21. 10.3389/fgene.2023.1168212

Powell, O., Gaynor, R.C., Gorjanc, G., Werner, C.R., & Hickey, J.M. (2020). A two-part strategy using genomic selection in hybrid crop breeding programs. bioRxiv, 1–46. 10.1101/2020.05.24.113258

R Development Core Team. (2022). R: A language and environment for statistical computing.

Sabadin, F., DoVale, J.C., Platten, J.D., & Fritsche-Neto, R. (2022). Optimizing self-pollinated crop breeding employing genomic selection: From schemes to updating training sets. Frontiers in Plant Science, 13. 10.3389/fpls.2022.935885

Schnell, F.W., & Utz, H.F. (1976). F1 Leistung und Elternwahl in der Zuchtung von Selbstbefruchtern. Ber Arbeitstag Arbeitsgem Saatzuchtleiter,

Toro, M.A., & Varona, L. (2010). A note on mate allocation for dominance handling in genomic selection. Genetics Selection Evolution, 42. 10.1186/1297-9686-42-33

Troyer, A.F., & Wellin, E.J. (2009). Heterosis Decreasing in Hybrids: Yield Test Inbreds. Crop Science, 49, 1969–1976. 10.2135/cropsci2009.04.0170

Utz, H.F., Bohn, M., & Melchinger, A.E. (2001). Predicting progeny means and variances of winter wheat crosses from phenotypic values of their parents. Crop Science, 41, 1470–1478

Varona, L., Legarra, A., Toro, M.A., & Vitezica, Z.G. (2018). Non-additive effects in genomic selection. Frontiers in Genetics, 9, 1–12. 10.3389/fgene.2018.00078

Wartha, C.A., & Lorenz, A.J. (2021). Implementation of genomic selection in public-sector plant breeding programs: Current status and opportunities. Crop Breeding and Applied Biotechnology, 21

Werner, C.R., Gaynor, R.C., Sargent, D.J., Lillo, A., Gorjanc, G., & Hickey, J.M. (2023). Genomic selection strategies for clonally propagated crops. Theoretical and Applied Genetics, 136, 1–17. 10.1007/s00122-023-04300-6

Wientjes, Y.C.J., Bijma, P., Van Den Heuvel, J., Zwaan, B.J., Vitezica, Z.G., & Calus, M.P.L. (2023). The long-term effects of genomic selection: 2. Changes in allele frequencies of causal loci and new mutations. Genetics, 225, iyad141

Wolfe, M.D., Chan, A.W., Kulakow, P., Rabbi, I., & Jannink, J.L. (2021). Genomic mating in outbred species: Predicting cross usefulness with additive and total genetic covariance matrices. Genetics, 220. 10.1093/genetics/iyab122

