## Supplemental Material S1 for "SimpleMating: R-package for Prediction and Optimization of Breeding Crosses Using Genomic Selection"

### 1. Background on usefulness estimation

The usefulness criterion (UC) was proposed by Schnell and Utz (1975), combining the concepts of genetic mean ($\mu$) and the genetic gain ($\Delta G$). It can be used to maximize the performance of the next generation. Its implementation and use as a criterion to parental cross selection has proven to return higher gains than the selection of combinations based on only truncation selection (Lehermeier et al., 2017; Allier et al., 2019). The following equation describes UC:

$$UC = \mu + \ ih\hat{\sigma}_g$$

where $\mu$ is the mean of a target cross (genetic mean), *i* represents the intensity of selection, $h$ is the squared root of the heritability (assumed to be 1), and $\hat{\sigma}_g$ represents the progeny genetic standard deviation within family for a target cross ($ih\hat{\sigma}_g$ term stands for genetic gain - $\Delta G$). In the equation above described, selection intensity ($i$) represents the mean of deviations from the population mean, and can be implemented as (Falconer & Mackay, 1996):

$$i = \Phi(x)/p,$$

$$\Phi(x) \ = \ \frac{1}{\sqrt{2\pi}} \ e^{-\frac{1}{2}x^2}$$

where $\Phi(x)$ is the probability density function of the standard normal distribution for *x*, $p$ is the selection proportion, and *x* is the mean deviation of the selected group as in:

$$x \ = \ \sum(y_i - \ \hat{y})/n$$

where $y_i$ is the $i^{th}$ selected observation, $\hat{y}$ is the population mean, and *n* refers to the number of selected observations. For the implementation in the analyses, we use the R formulae, as follows (Mackey, 2020):

$$i = qnorm(dnorm(1-p))/p,$$

where *qnorm* is the quantile normal distribution and *dnorm* is the density of the normal distribution. It returns to us the selection intensity for any level of selection proportion ($p$).

The estimation of cross mean ($\mu$) and cross variance ($\hat{\sigma}_g$), for progeny standard deviation) to compose the usefulness criterion for a set of crosses between candidates to parents is discussed below. For such end, a breeding population is assumed, likewise an allele dosage matrix, estimates of marker effects, and a genetic map or a linkage disequilibrium matrix for calculate a recombination matrix for the target parent population, using bi-parental crosses.

**2. Prediction of cross mean**

The genetic mean of a future specific cross can be estimated by considering only additive effects (estimated breeding value – *ebv*), where the genetic mean of a progeny is computed as the mean parental average ($\mu_{ebv}$). In addition, in traits with additive and dominance gene actions, we can capitalize the heterosis in the F1 population by considering the dominance effects in the equation, where the cross total genetic value (*tgv*) is estimated for a cross, by considering alleles frequencies and the additive/dominance effects of markers for a target trait ($\mu_{tgv}$). The following equations are used for the estimation (Falconer & Mackay, 1996):

$$\mu_{ebv} = \frac{GEBV_{P1} + GEBV_{P2}}{2}$$

$$\mu_{tgv} = \sum_{k=1}^{p} a_k\left(p_{ik} + \mathrm{q}_{ik} + \mathrm{y}_k\right) + \mathrm{d}_k\left[2\mathrm{p}_{ik}\mathrm{q}_{ik} + \mathrm{y}_k\left(\mathrm{p}_{ik} - \mathrm{q}_{ik}\right)\right]$$

where *GEBV* represents the genomic *ebv* for each parent, *a* and *d* represents the additive and dominance effects of the markers, respectively, *p* and *q* represent the allele frequency for a marker, and $y$ represents the difference in allele frequency between individuals involved in a cross. Both descriptions ($\mu_{ebv}$ and $\mu_{tgv}$) are implemented and can be explicitly estimated using the functions `getMPA()` and `getTGV()`, respectively. In addition, they are internally estimated in all usefulness functions (`getUsefA` and `getUsefAD`, respectively).

The index-based mean for more than one trait ($\mu t$) for a given cross is possible for both cases above described. The same formulas are used to predict the performance of each trait individually, and an index-based trait is estimated by weighting each cross mean by an economic weight set by accordingly, as follows:

$$\mu t_{ebv} = v' \mu_{ebv}$$

$$\mu t_{tgv} = v' \mu_{tgv}$$

where $v$ is a vector of $1 \times t$ ($t$ represents the number of traits) and $\mu_{ebv}$ and $\mu_{tgv}$ represents the mean estimated for each cross (before defined), with dimensions $n \times t$ (where $n$ represents the total number of crosses). Both scenarios are implemented in the functions `getMAP()` and `getTGV()`, and weights for each trait should be given, while present internally in the usefulness functions `getUsefA_mt` and `getUsefA_mt`.

### 3. Prediction of cross variance

In a similar way than the prediction of the cross mean, the additive progeny variance ($\hat{\sigma}^2_{pa}$) and dominance progeny variance ($\hat{\sigma}^2_{pd}$) of a cross were implemented. In a case of pure additivity, the additive progeny variance for a target cross is (Lynch & Walsh, 1988; Falconer & Mackay, 1996; Bonk et al., 2016; Lehermeier et al., 2017):

$$\hat{\sigma}^2_{pa} = a' \Omega a$$

where $a$ represents the vector of markers' additive effect, and Ω is the covariance of the progeny for the target cross ($j \times j$, where $j$ is the number of markers for the population). Considering fully inbred lines (*e.g.*, DH or RILs), the Ω component is (Lehermeier et al. (2017)):

$$\Omega = \begin{pmatrix} 4\pi_j(1-\pi_j) & \cdots & 4D_{jl} \\ \vdots & \ddots & \vdots \\ 4D_{lj} & \cdots & 4\pi_l(1-\pi_l) \end{pmatrix}$$

where $\pi_j/\pi_l$ is the allele frequency for a marker *j*/*l* and $D_{jl}$ the disequilibrium parameter between markers $j$ and $l$. The diagonal values (variance) hold from the equation: $(1 + F)2\pi_j(1 - \pi_j)$, where F is the inbreeding coefficient (considering F = 1, *i.e.*, DH lines are fully inbred). This equation implies that $\pi_j \in \{0, ½, 1\}$, while for a DH line, the diagonal of Ω matrix is either 0 (for non-segregating parents) and 1 (for parents differing at the locus) (Lehermeier et al., 2017). In this same sense, the off-diagonal for DH case, is $4D_{jl}$, being $D_{jl}$ a proxy of the disequilibrium parameter among parental lines (noted as $D_{jl}^*$) and the recombination frequency expected in the parental population ($c_{jl}$), as follows: $D_{jl} = (1 - 2c_{jl})D_{jl}^*$. Moreover, it also holds that $D_{jl} \in \{0, ¼, -¼\}$, assuming the same allele at one of both loci, two loci in coupling phased, and two loci in repulsion phase, respectively (Lehermeier et al., 2017). In this sense, the additive progeny variance for DH lines from F1 population can be estimated as: $\Omega = 4D_{jl}^*(1 - 2c_{jl})$. The same authors extended the estimation of progeny variance for cases including different generations for DH derivation and recombinant inbred lines or RILs (Table 1). All four formulas are implemented in **SimpleMating** for the estimation of cross variance.

Table 1. For a pure additive trait, the formulas implemented in SimpleMating for covariance (Ω) estimation. Adapted from Lehermeier et al. (2017). DH = doubled haploids. RIL = recombinant inbred lines. $D_{jl}^{*}$ = linkage disequilibrium in the parental lines. $c_{jl}$ = recombination expected in the base population. k = generation.

| Parental population | Progeny covariance estimation (Ω) |
|---|---|
| ***DH (F1)*** | $4D_{jl}^{*}(1-2c_{jl})$ |
| $\boldsymbol{DH^{k}}$ | $4D_{jl}^{*}(\Sigma_{r=1}^{k}(0.5(1-2c_{jl}))^{r} + (0.5(1-2c_{jl}))^{k})$ |
| $\boldsymbol{RIL^{k*}}$ | $4D_{jl}^{*}(\Sigma_{r=1}^{k}(0.5(1-2c_{jl}))^{r})$ |
| $\boldsymbol{DH/RIL^{\infty}}$ | $4D_{jl}^{*}(1-4c_{jl})/(1+2c_{jl})$ |

* for RILs, k=1 represents F2 population.

The recombination frequency expected in the parental population ($c_{jl}$) is based on linkage disequilibrium (LD) among gametes from the population of candidates to parents (Lehermeier et al. (2017), Bonk et al. (2016)). It depends on the map distance between two marker loci ($x$), easily estimated via Haldane's mapping function using the genetic map information (Haldane, 1919), as implemented in **SimpleMating**, as following:

$$c_{jl} = -\frac{1}{2}ln(1-2x)$$

Aside pure additive control, progeny variance estimation accounting for additive and dominance trait effects was also implemented in **SimpleMating**. The total genetic progeny variance ($\hat{\sigma}_{gp}^{2}$) (that accounts for both, additive and dominance effects) is as following:

$$\hat{\sigma}_{gp}^{2} = \hat{\sigma}_{pa}^{2} + \hat{\sigma}_{pd}^{2}$$

In such case, we used the phased diplotypes from the candidates to parents for the estimation of covariance parameter (Ω). For additive and dominance progeny variances, the following holds (Allier et al., 2019; Wolfe et al., 2021):

$$\hat{\sigma}_{pa}^2 = a'\Omega a$$

$$\hat{\sigma}_{pd}^2 = d'\Omega^2 d$$

where, $d$ is the vector of additive effects for each marker. Furthermore, the covariance $\Omega$ matrix for the genotypes is estimated from the haplotypes for each parent ($D_p$) (Bonk et al., 2016):

$$D_p = (1 - 2c_{jl}) \odot D_{haplo}$$

where $D_{haplo} = \frac{1}{2} H'H - pp'$, being $H$ the haplotype information for each parent and $p$ the allele frequency matrix for the individual at each locus. Hence, $\Omega = D_{p1} + D_{p2}$. For the dominance effect, $\Omega^2$ has the effect of squaring all elements (Lynch & Walsh, 1988).

As before noticed, the disequilibrium parameter between pair of parents ($D_{jl}$) rely on the knowledge of phased haplotypes, for the case where additive and additive plus dominance gene actions controls the trait (largely used in outbreed species). One alternative implemented in SimpleMating is to calculate the variance matrix between pair of crosses via unphased haplotypes. The formula still accounts for the recombination matrix ($c_{jl}$) for the group of parents and a variance matrix was estimated between pair of parents $H_{jj}^*$, as follows:

$$\delta = \left(1 - 2c_{jl}\right)$$

$$\Omega = (\delta - \mathrm{diag}(\delta)) + H_{jj}^*$$

$H_{jj}^*$ is a diagonal matrix of length $j$ x $j$ (number of markers) with elements for additive effects:

$$H_{jj}^* = \begin{cases} 0, & if\ both\ parents\ are\ homozygous \\ 0.5, & if\ one\ is\ homozygous\ and\ the\ other\ one\ is\ heterozigous \\ 1, & if\ both\ parents\ are\ heterozigous \end{cases}$$

and for dominance values, the same hold for the $H^*_{jj}$, being the approximation for the diagonal as follows:

$$H^*_{jj} = \begin{cases} 0, & \textit{if both parents are homozygous} \\ 1, & \textit{if at least one parent is heterozigous} \end{cases}$$

In addition, the matrix $C_{jl}$ is essentially derived from a genetic map. A genetic map is based on the recombination frequency between markers for a target population. For SimpleMating optimization, the genetic map can be used. With those values, using the Haldane function (Haldane 1919), we can estimate the recombination for each pair of markers in the data. Another possibility is to obtain the linkage disequilibrium matrix for the markers and use this information to get $C_{jl}$, as in $C_{jl} = 1 - LD$. We implemented that in all usefulness functions in SimpleMating.

When we investigate a multi-trait framework, the variance estimated for several traits follows the construction of covariance matrix at trait-level with the dimension of the number of traits analyzed ($t \times t$). For such, we build a matrix of covariances at trait level (T) by calculating in the first step every trait variance and the covariances between each pair of traits, using the estimation of covariance parameter (Ω) such before stated, as follows (Bonk et al., 2016):

$$\hat{\sigma}^2_{T1} = \alpha_{T1}{}'\Omega\alpha_{T1},$$

$$\hat{\sigma}^2_{T2} = \alpha_{T2}{}'\Omega\alpha_{T2},$$

$$\hat{\sigma}^2_{T1T2} = \alpha_{T1}{}'\Omega\alpha_{T2},$$

then,

$$T = \begin{bmatrix} \hat{\sigma}^2_{T1} & \hat{\sigma}^2_{T1T2} \\ \hat{\sigma}^2_{T1T2} & \hat{\sigma}^2_{T2} \end{bmatrix},$$

where the T matrix is used in the following equation for the estimation of a cross variance for a multi-trait framework ($\hat{\sigma}^2_{SI}$) using the weights for each trait ($v$):

$$\hat{\sigma}^2_{SI} = v'Tv$$

### References


Allier, A., Moreau, L., Charcosset, A., Teyssèdre, S., & Lehermeier, C. (2019). Usefulness criterion and post-selection parental contributions in multi-parental crosses: Application to polygenic trait introgression. *G3: Genes, Genomes, Genetics*, *9*, 1469–1479. https://doi.org/10.1534/g3.119.400129

Bonk, S., Reichelt, M., Teuscher, F., Segelke, D., & Reinsch, N. (2016). Mendelian sampling covariability of marker effects and genetic values. *Genetics Selection Evolution*, *48*, 1–11. https://doi.org/10.1186/s12711-016-0214-0

Falconer, D.S., & Mackay, T.F.C. (1996). Introduction to quantitative genetics. *Harlow, Essex, UK: Longmans Green*, *3*, 280

Haldane, J.B.S. (1919). The combination of linkage values and the calculation of distances between the loci of linked factors. *J Genet*, *8*, 299–309

Lehermeier, C., Teyssèdre, S., & Schön, C.C. (2017). Genetic gain increases by applying the usefulness criterion with improved variance prediction in selection of crosses. *Genetics*, *207*, 1651–1661. https://doi.org/10.1534/genetics.117.300403

Lynch, M., & Walsh, B. (1988). *Genetics and Analysis of Quantitative Traits*. Sinauer, Sunderland, MA.

Mackey, I. (2020). Selection Intensity

Wolfe, M.D., Chan, A.W., Kulakow, P., Rabbi, I., & Jannink, J.L. (2021). Genomic mating in outbred species: Predicting cross usefulness with additive and total genetic covariance matrices. *Genetics*, *220*. https://doi.org/10.1093/genetics/iyab122
