## Supplemental Material S2 for "SimpleMating: R-package for Prediction and Optimization of Breeding Crosses Using Genomic Selection"

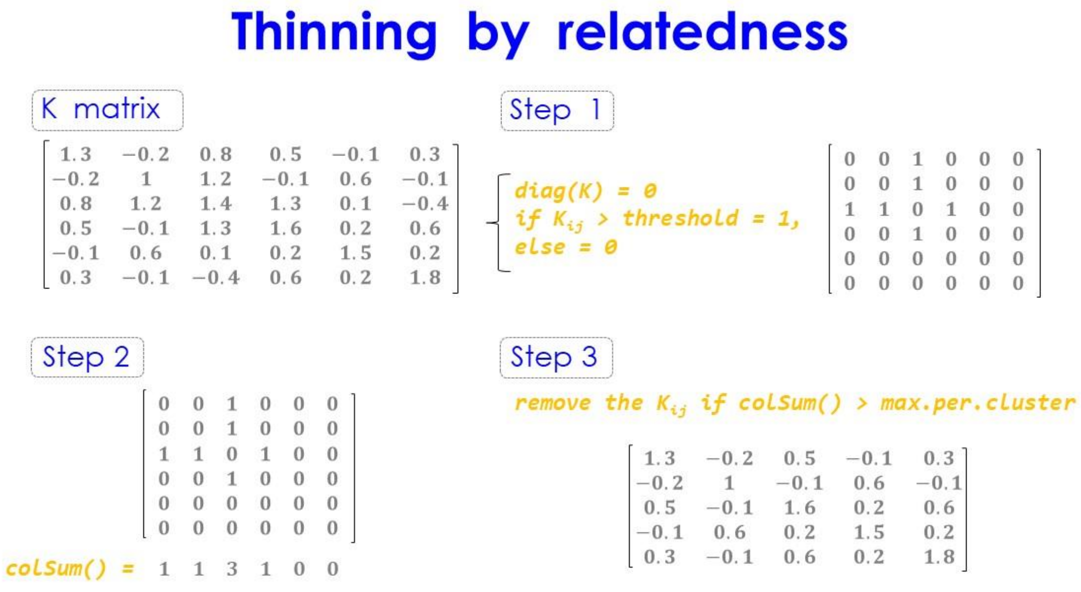


**Figure 1 - Simplified representation of the function '`relateThinning`()' from the SimpleMating.** The relationship matrix K (pedigree- or marker-based) is coded in a matrix with 0 and 1 based on the level of relatedness of a target genotype with the others, using the given *threshold*. Every value above the threshold receives the number 1, otherwise 0 (Step 1). After, all columns are summed (Step 2) and after one iteration per row number, we remove the genotype with sum higher than *max.per.cluster* limit (Step 3), based on the value set via criterion argument.

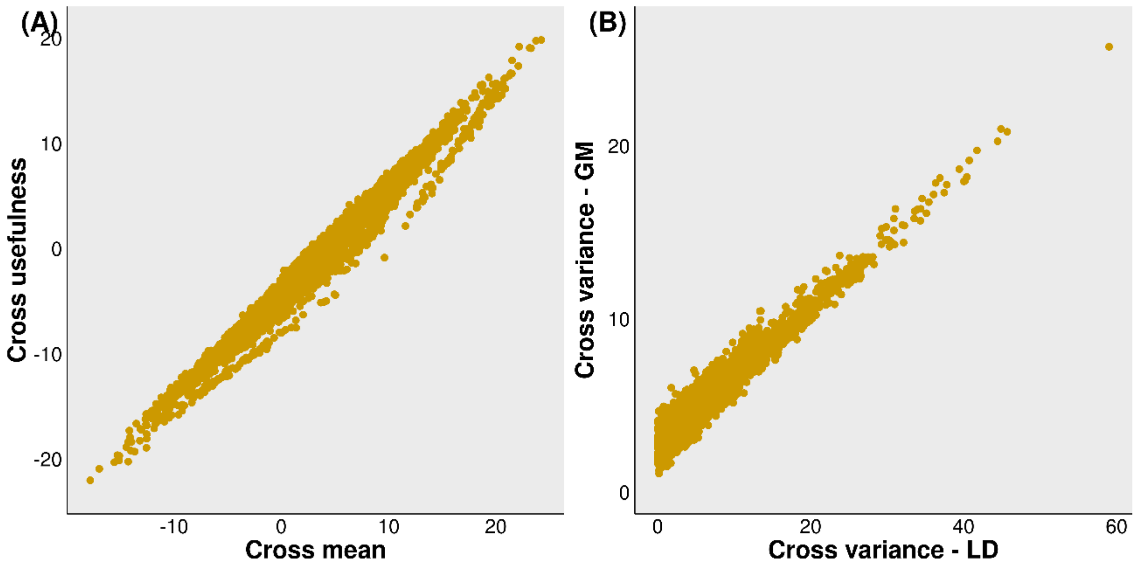


**Figure 2 – Correlation plots from the outcomes of the usefulness function (*getUsefA*).** A) Cross usefulness by cross mean and B) cross variance estimated using the recombination matrix estimated from a genetic map (GM, Y-axis) and a linkage disequilibrium matrix (LD, X-axis). A total of 4950 crosses were predicted from the example above mentioned. Every single dot represents one biparental cross.
